# Immune Correlates of Hyperglycemia and Vaccination in a Non-human Primate Model of Long-COVID

**DOI:** 10.1101/2023.09.22.559019

**Authors:** Clovis S Palmer, Chrysostomos Perdios, Mohamed Abdel-Mohsen, Joseph Mudd, Prasun Datta, Nicholas J. Maness, Gabrielle Lehmicke, Nadia Golden, Lihn Hellmer, Carol Coyne, Kristyn Moore Green, Cecily Midkiff, Kelsey Williams, Rafael Tiburcio, Marissa Fahlberg, Kyndal Boykin, Carys Kenway, Kasi Russell-Lodrigue, Angela Birnbaum, Rudolf Bohm, Robert Blair, Jason Dufour, Tracy Fischer, Ahmed A Saied, Jay Rappaport

**Affiliations:** Tulane National Primate Research Center, Covington, LA, USA; Department of Microbiology and Immunology, Tulane University School of Medicine, New Orleans, LA, USA; The Wistar Institute, Philadelphia, PA, USA; Division of Experimental Medicine, Department of Medicine, University of California, San Francisco; Department of Pathology and Laboratory Medicine, Tulane University School of Medicine

**Keywords:** SARS-CoV-2, Diabetes, Long-Covid, PACS, Vaccine, metabolism, immunometabolism

## Abstract

Hyperglycemia, and exacerbation of pre-existing deficits in glucose metabolism, are major manifestations of the post-acute sequelae of SARS-CoV-2 (PASC). Our understanding of lasting glucometabolic disruptions after acute COVID-19 remains unclear due to the lack of animal models for metabolic PASC. Here, we report a non-human primate model of metabolic PASC using SARS-CoV-2 infected African green monkeys (AGMs). Using this model, we have identified a dysregulated chemokine signature and hypersensitive T cell population during acute COVID-19 that correlates with elevated and persistent hyperglycemia four months post-infection. This persistent hyperglycemia correlates with elevated hepatic glycogen, but there was no evidence of long-term SARS-CoV-2 replication in the liver and pancreas. Finally, we report a favorable glycemic effect of the SARS-CoV-2 mRNA vaccine, administered on day 4 post-infection. Together, these data suggest that the AGM metabolic PASC model exhibits important similarities to human metabolic PASC and can be utilized to assess therapeutic candidates to combat this syndrome.

## 1. Introduction

Between 10-30% of people infected with SARS-CoV-2 develop long-term health complication, named post-acute sequelae of SARS-CoV-2 (PASC) or Long-COVID ^1–4^. Metabolic diseases, including type 2 diabetes (T2D)^5^, as well as conditions with less obvious metabolic undertones such as myalgic encephalomyelitis/chronic fatigue syndrome (ME/CFS), thrombosis and neuropsychiatric sequelae (brain fog) embody the broad spectrum of long-COVID symptoms or PASC ^1, 6–10^. In fact, a 30-50% elevated T2D incidence following acute resolution of SARS-CoV-2 infection has been reported^11^. Several lines of evidence suggest a hyperinflammatory response against SARS-CoV-2 as being critical to the severity of acute COVID-19^12^, as well as development of metabolic PASC such as hyperglycaemia^3, 11^, metabolic associated fatty liver disease (MAFLD)^13^, and cardiovascular diseases (CVD)^14, 15^.

An existing paradigm postulates that the balance between virus survival and effective host responses is based on metabolic reprogramming of nutrients, primarily in immune cells^16^. However, the extent to which early disruptions in systemic immune and metabolic homeostasis contribute to the evolution and symptoms of metabolic PASC remains unclear. While pre-existing diabetes has been linked to more severe COVID-19 outcomes and higher mortality^17, 18^, new-onset hyperglycemia and diabetic ketoacidosis is also observed in SARS-CoV-2 infected individuals with no prior evidence of diabetes and have been associated with poor COVID-19 outcomes^19, 20^.

Glucose homeostasis is maintained largely by hormonally-regulated glucose uptake by tissues, such as the liver^21, 22^, as well as the gut microbiota^23^. However, hepatic glucose production via gluconeogenesis and glycogenolysis represents additional glucometabolic checkpoints^24^. In addition, β-cell dysfunctions, potentially due to early pancreatic infection by SARS-CoV-2, may partially contribute to altered glucose homeostasis^25–27^. Increased levels of circulating inflammatory molecules, including chemokines, have been shown to directly associate with impaired glucose homeostasis in non-infectious diseases^28^. This includes CCL25, a cytokine known to impair pancreatin β-cell insulin secretion and is capable of inducing proinflammatory cytokine responses^29^. Thus, a preponderance of evidence supports a key role for inflammation in the pathogenesis of hyperglycemia and T2D^30–32^. However, the mechanisms by which SARS-CoV-2 infection promotes prolonged hyperglycemia are poorly understood due to the lack of appropriate animal models for metabolic PASC. Here, we developed such a model and used it to interrogate potential mechanism that underlie the development of metabolic PASC. We also investigated whether administration of the BNT162b2 (Pfizer/BioNTech) vaccine during acute SARS-CoV-2 infection could ameliorate immunometabolic dysregulation.

## 2. Results

### 2.1. Study groups

African green monkeys (AGMs; *Chlorocebus aethiops sabaeus*) were followed up weekly for 18 weeks with complete virologic, physical, clinical assessments, blood chemistry and immunometabolic profiling. Some assessments were conducted biweekly after 4 weeks (Fig. 1a). One female (PB24), age 19.32 years (6.40 kg) was sent for necropsy at week 8 due to anorexia. There were no significant differences in the median age and weight between the vaccinated and unvaccinated group (see Table 1 for detailed group demographics).

**Figure 1.**
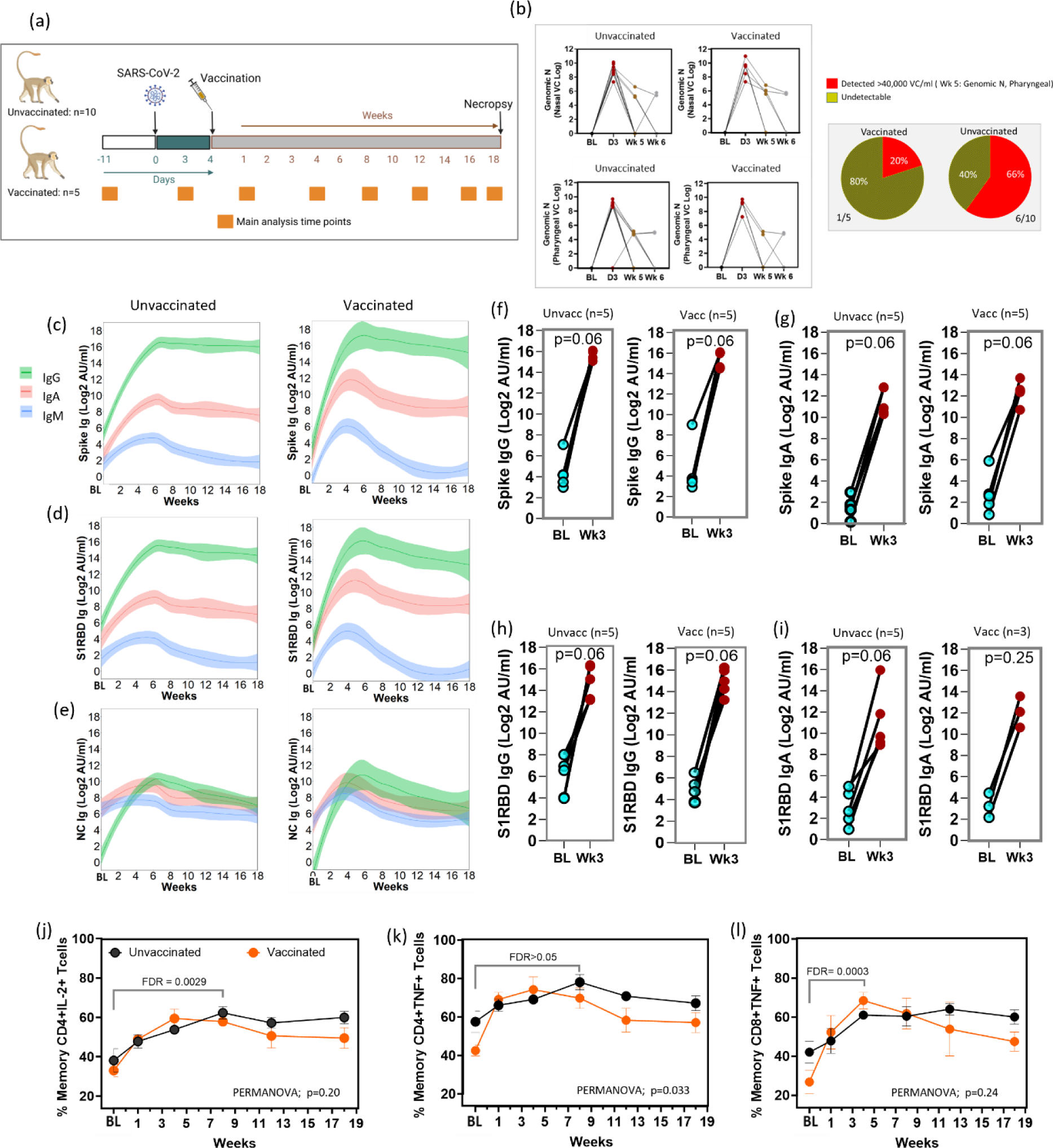
SARS-CoV-2 infection of AGM is associated with a strong virologic, immunologic and hypersensitive response. (a) An overview of the study design and major analysis timepoints. (b) Kinetics of nasal and pharyngeal SARS-CoV-2 genomic nucleocapsid (N) RNA in unvaccinated and vaccinated animals (left), and pie charts showing the percentage of animals with detectable pharyngeal genomic RNA in the vaccinated vs unvaccinated group at week 5 (right). (c) Temporal assessment of antibody responses against spike, (d) S1RBD, and (e) nucleocapsid proteins in unvaccinated and vaccinated animals. (f) Comparative time point analysis (Baseline, BL, vs week, Wk, 3) of IgG, and (g) IgA responses against Spike protein. (h) Comparative time point analysis (Baseline, BL, vs week, Wk, 3) of IgG, and (i) IgA responses against S1RBD protein in unvaccinated and vaccinated animals. (j) The percentage of memory CD4+ T cells expressing IL-2, and (k) TNFα over time in response to PMA/ionomycin stimulation. (l) The percentage of memory CD8+ T cells expressing TNFα over time in response to PMA/ionomycin stimulation. Wilcoxon matched pairs signed rank test with Benjamini-Hochberg correction (FDR) was used to compare time points. PERMANOVA was used for temporal comparisons between the unvaccinated and the vaccinated group. Error bars represent SEM. BL: 1.5 weeks pre-infection.

**Table 1.** Characteristics of the study population, and the vaccinated and unvaccinated subpopulations.

| Characteristics | Study population | Unvaccinated | Vaccinated (Pfizer) | *p* |
| --- | --- | --- | --- | --- |
| **N** | 15 | 10 | 5 |  |
| **Sex (F/M)** | 13/2 | 9/1 | 4/1 |  |
| **Age (years), median (min-max)** | 8.970<br>(7.92 - 19.32) | 8.97<br>(7.92 -17.92) | 8.97<br>(7.92-19.32) | 0.53 |
| **Weight (Kg) (min-max)** | 4.35<br>(3.75-6.55) | 4.35<br>(3.75-5.45) | 4.60<br>(4.15-6.55) | 0.53 |

### 2.2. Dynamics of SARS-CoV-2 over time

To confirm infection and study viral dynamics we assessed sub genomic viral RNA (sgRNA), a correlate of actively replicating virus, and genomic RNA (gRNA) levels by quantitative real-time PCR in nasal and pharyngeal swabs. All animals had detectable sgRNA (>3.64 × 10^6^) at day 3 in both nasal and pharyngeal swabs. At week 5, only 2 animals had detectable sgRNA (>1.15 × 10^5^) in nasal swabs, and no detection in pharyngeal swabs. At day 3, all animals had detectable gRNA (> 2.05 × 10^7^) in nasal swabs, while 14 had detectable gRNA (>1.71 × 10^6^) in pharyngeal swabs. At week 5, seven animals had detectable gRNA (1.47 × 10^5^ - 6.53 × 10^6^) in nasal swabs, and seven detectable (4.90 × 10^4^ - 1.51 × 10^5^) in pharyngeal swabs. The kinetics of gRNA were similar in vaccinated and unvaccinated, with virus peaking at day 3 and substantially declining by week 5 (Fig. 1b). At week 5, 20% and 66% of the vaccinated and unvaccinated animals respectively had detectable gRNA in pharyngeal swabs (Fig. 1b).

### 2.3. Changes in immune compartments, antibody, and hypersensitive responses during long-term follow-up

To evaluate changes in systemic immune cell compartments over time, we assessed major immune subsets which have previously been implicated in the severity of acute COVID-19 disease. On day 3 post infection (p.i.), we observed a significant increase in monocyte percentage (p=0.0004) and absolute number (p=0.002) in infected animals, returning to baseline levels by week two, followed by a modest decline up to about week 7 (Fig. S1. a-b). The percentage and absolute counts of lymphocytes and neutrophils remained relatively constant (Fig. S1 c-f) in both the vaccinated and unvaccinated groups. The declining levels of circulating monocytes post-acute infection may signify potential infiltration of these cells into tissues, consistent with elevated tissue inflammation during acute SARS-CoV-2 infection.

We observed a steep induction of IgG and IgA responses against SARS-CoV-2 Spike and S1RBD proteins, peaking between 3-6 weeks (Fig. 1c-d), and maintained elevated up to 18 weeks p.i. in both the vaccinated and unvaccinated groups. There was also an appreciable IgA and IgG response against SARS-CoV-2 nucleocapsid, with the IgG levels remaining elevated up to 15 weeks (Fig. 1e). The early IgM response against SARS-CoV-2 was noticeable but of a lesser magnitude than the IgA and IgG responses, and sharply declined towards baseline after 5-week p.i. (Fig. 1c-e). Although not reaching statistical significance, cumulatively there was a substantially elevated IgG and IgA antibody responses against the Spike (Fig. 1f-g) and S1RBD (Fig. 1h-i) proteins at week 3 p.i. relative to baseline (1.5 weeks pre-infection). These results closely reflect the kinetics and preferential induction of anti-SARS-CoV-2 IgA, IgG, and IgM responses in infected humans^33^.

There were no significant differences in the magnitude of antibody responses between the vaccinated and unvaccinated groups over time (PERMANOVA; p>0.05, data not shown). However, there was a slight trend towards a higher induction of IgA response towards the spike protein, as well as IgG response towards nucleocapsid protein (PERMANOVA; p>0.05, data not shown).

T-cell inflammatory response to polyclonal stimulation through phorbol 12-myristate 13-acetate (PMA) and the calcium ionophore ionomycin (I) has been reported in severe COVID-19 cases^12^. To examine potential similarities, we exposed longitudinally collected PBMCs from SARS-CoV-2 infected AGMs to PMA/I for 6 hours and examined activation markers and intracellular levels of cytokines in T cells subsets. Naïve and memory populations were defined using CD95 and CD28 markers according to established gating strategies (Fig. S3). Cumulatively, across groups, there was a significantly increased percentage of memory CD4+ T cells expressing IL-2 (Fig. 1j; FDR< 0.01) between baseline and all following weeks p.i. Cumulatively, across timepoints, there was a significantly lower percentage of memory CD4+ T cells expressing TNF in the vaccinated group versus the unvaccinated one (Fig. 1k; PERMANOVA; p=0.033). Except for week 1 (FDR> 0.05), cumulatively, the percentage of memory CD8+ T cells expressing TNF was significantly increased relative to baseline (Fig. 1l; FDR< 0.01). Taken together, these data indicate that the AGM SARS-CoV-2 infection model mirrors several immunologic similarities to those reported in SARS-CoV-2 infected humans.

### 2.4. Vaccination post SARS-CoV-2 infection improves long-term glycemic control

Early metabolic changes during SARS-CoV-2 infection are likely to influence long-term manifestations of COVID-19. We have previously shown in AGMs that SARS-CoV-2 infects pancreatic ductal, and endothelial cells and associates with new-onset hyperglycemia^34^, an observation that recapitulates findings in humans, supporting pancreatic tropism of SARS-CoV-2^26, 27^. SARS-CoV-2 infection may also promote hyperglycemia by inducing excess hepatic glucose production through gluconeogenesis or glycogenolysis independent of pancreatic function or insulin effects^35^. We therefore analyzed serum glucose levels over time and found it to be significantly elevated (n=15, mean: 102.2 mg/dL, range: 76 −154, p<0.0001) as early as three days post-infection relative to the latest baseline values (n=15, mean: 71.1 mg/dL, range: 51-86; Fig. 2a). This elevated blood glucose is well above the normal range independently reported for male (mean: 81.7 ± 18.7 mg/dL) and female (mean: 80.3 ± 18.7 mg/dL) AGMs of Caribbean origin^36^. Composite longitudinal analysis showed significantly higher blood glucose in the unvaccinated group (PERMANOVA; p=0.001) than the vaccinated group over time. This is supported by a higher percentage of animals with glucose levels above 100 mg/dL in the unvaccinated group (Fig. 2b). Moreover, a statistically significant persistence of hyperglycemia was maintained up to 12 weeks p.i. in the unvaccinated group (Fig. 2c). We excluded glucose reading for weeks 10 and 18 due to in-house clinical procedures likely to impact transient blood levels, namely the movement of animals from BSL3 to BSL2 (week 10) and return to BSL3 at week 18. There was a modest positive but non statistically significant relationship between day 3 nasal viral sgRNA and gRNA, and blood glucose levels at week 17 across groups (Fig. 2d). Peak IgA response (week 3) against nucleocapsid and S1RBD was lower in unvaccinated SARS-CoV-2 infected animals with higher preceding blood glucose (week 2; Fig. 2e).

**Figure 2.**
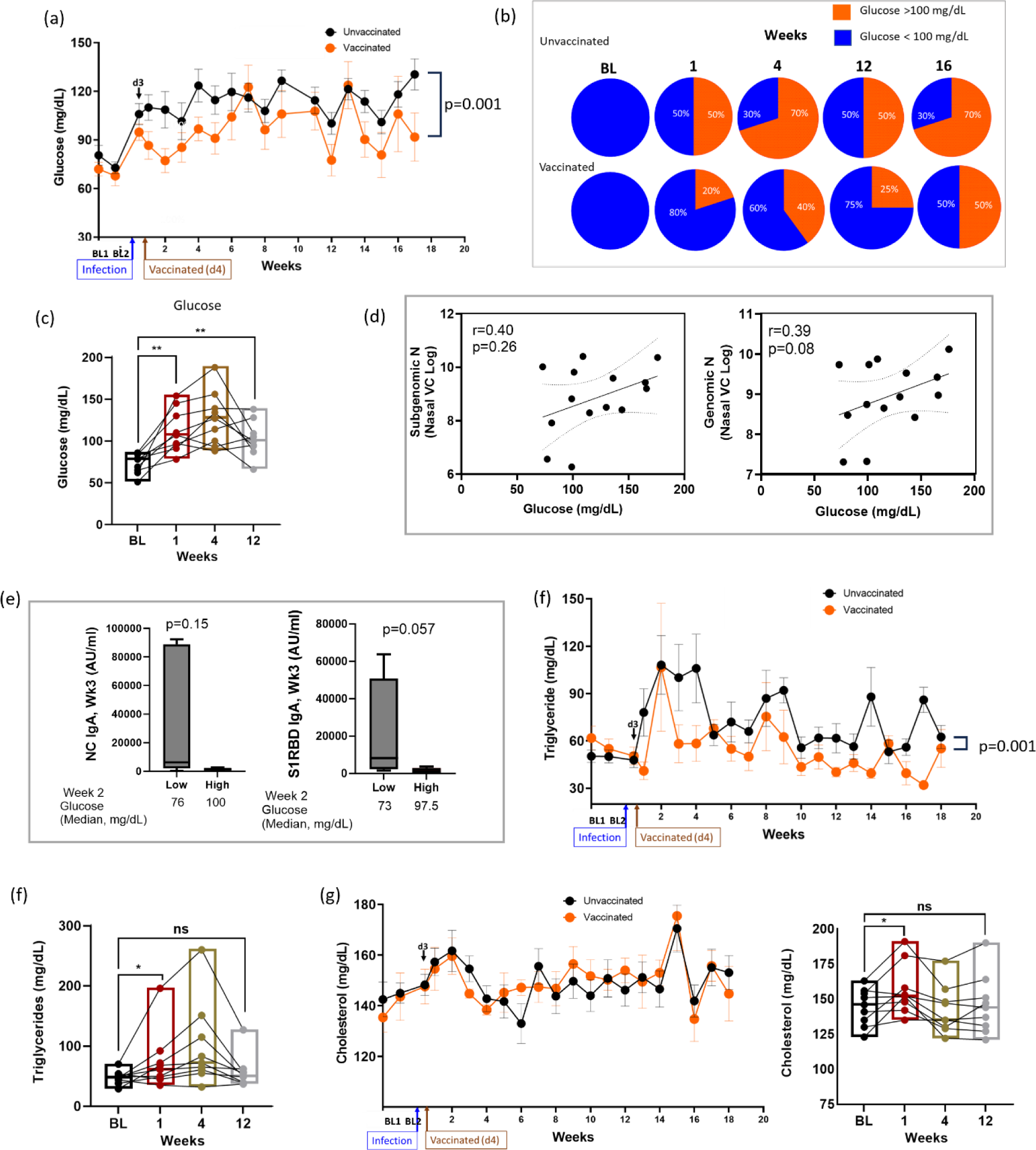
SARS-CoV-2 infection of AGM is associated with elevation and persistence of blood glucose concentration even after undetectable virus. (a) Levels of plasma glucose following infection over time. PERMANOVA analysis was used for statistical analysis to examine the cumulative difference between glucose levels in the two groups over time. (b) Pie charts showing the proportion of animals with glucose >100 mg/dL at baseline (BL) and week 1, 4, 12 and 16. (c) Glucose levels at BL and remained so at week 12 post-infection. Analysis conducted using Wilcoxon matched pairs signed rank test. (d) Pearson correlation between glucose levels and sub genomic nucleocapsid (N) and genomic N at day 3 post infection. (e) IgA response against N and S1RBD at week (Wk) 3 against unvaccinated animals with low and high glucose in the preceding week (week 2). (f) Triglyceride levels over time between to the vaccinated and unvaccinated group. PERMANOVA was used for statistical analysis (g) Total cholesterol levels between the unvaccinated and vaccinated group. Key: BL1 = baseline 1 (6.5 weeks pre-infection); BL2 = baseline 2 (1.5 weeks pre-infection); d3 = day 3; ns = non-significant; * = p<0.05; ** = p< 0.01. Error bars represent SEM.

Although plasma triglyceride levels peaked around week 2, the levels were already significantly elevated by week 1 in the unvaccinated group (Fig. 2f). Cumulatively, triglyceride levels were significantly higher in the unvaccinated group over time (PERMANOVA; p=0.001). However, this elevation was not maintained consistently beyond baseline over the study period (Fig. 2f). We recorded no significant difference in cholesterol levels between the two groups over time (Fig. 2g). Taken together, we reveal a significant induction and persistent hyperglycemia in SARS-CoV-2 infected AGMs, suggesting this experimental design as a potential model of metabolic PASC. Furthermore, we found that vaccination during acute infection could have a positive effect on glycemic control.

### 2.5 Induction of early inflammatory disturbances in AGM long-COVID model

We next undertook an unbiased approach to identify systemic cellular processes associated with SARS-CoV-2 infection in AGMs. We utilized the OLINK Proximity Extension Assay (PEA) technology, which has exceptional specificity requiring the binding of two matched-paired antibodies tagged with a unique DNA sequence followed by PCR amplification and signal generation. We used a Target 96 Inflammation Panel assessing proteins associated predominantly with apoptosis, immune activation and inflammatory responses, MAPK cascade, chemotaxis, and chemokine secretion. Sixty-five analytes were analyzed following stringent data QC of which 14 were significantly elevated, and one (IL-8) significantly reduced at week 1 p.i. (Fig. 3a, Table S1). Principal component analysis of these differentially regulated analytes expectedly showed a marked separation between baseline and week 1, corroborating the heatmap analysis (Fig. 3a, Table S1). The 14 elevated analytes include some with known chemotaxis and other inflammatory processes. Five analytes, C-C Motif Chemokine Ligand 25 (CCL25), CUB domain-containing protein 1 (CDCP1), FMS-like tyrosine kinase 3 ligand (Flt3L), C-C Motif Chemokine Ligand 8 (CCL8) and Stem Cell Factor (SCF) maintained significance (FDR <0.05) following Benjamini–Hochberg (BH) correction (Table insert, Fig. 3a). We highlight representative chemokines (CCL25, CCL-8, CCL19) and inflammatory molecules (IL-18, TNF) showing significant elevation at week 1, normalizing by week 12 (Fig. 3b-c).

**Figure 3.**
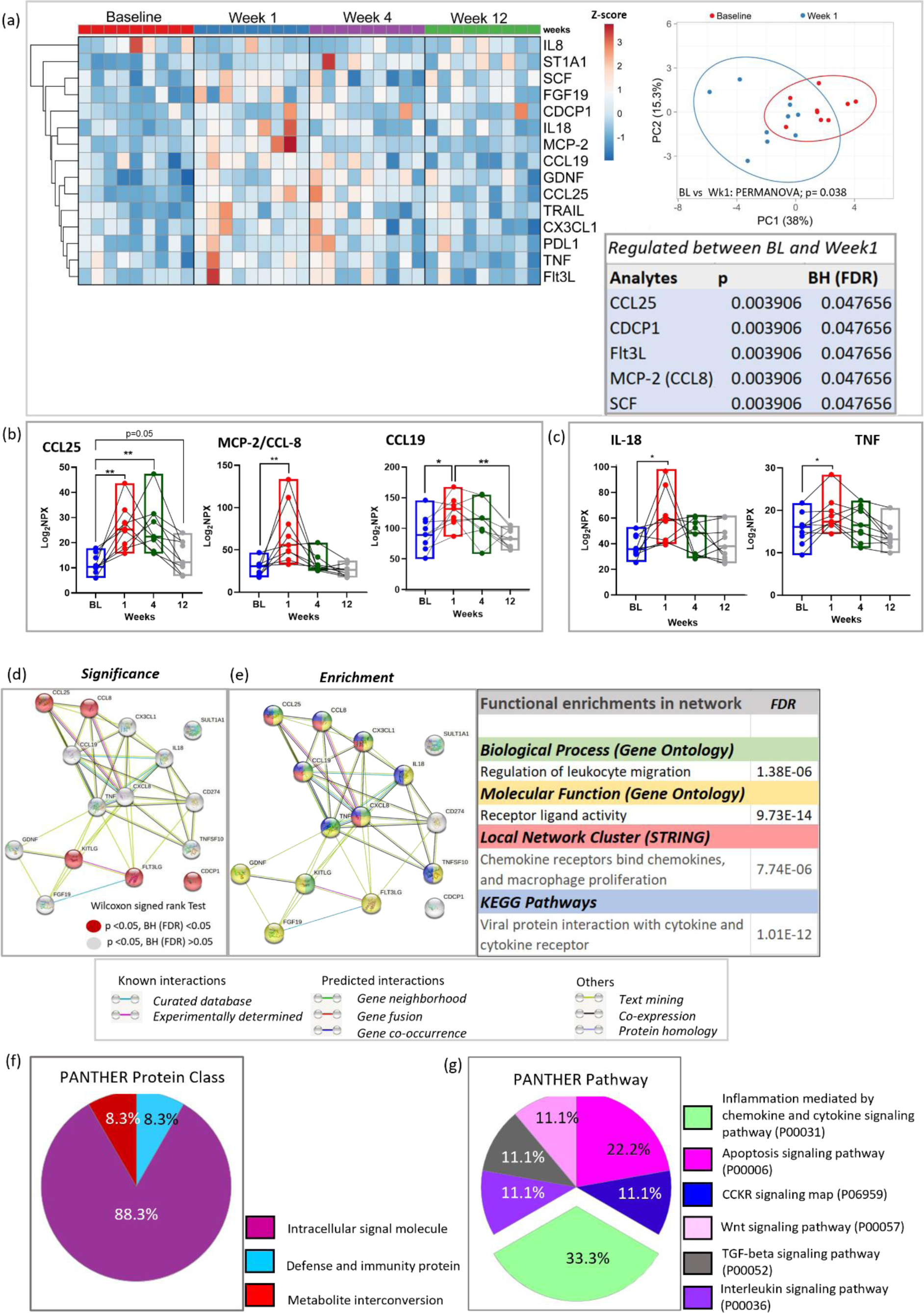
Elevated inflammatory mediators during acute SARS-CoV-2 infection of AGM. (a) Heat-map depicts 15 plasma analytes dysregulated between baseline and week 1 during SARS-CoV-2 infection (p<0.05). Heat colors show standardized Z-scores; red indicates highest levels of analytes, and blue indicates lowest levels. Statistical significance was determined using the Wilcoxon signed rank test. PCA plot confirms significant differences in analyte expression at baseline (red) and week 1 (blue). Box insert shows 5 dysregulated analytes remaining significantly regulated after Benjamini-Hochberg (BH) correction (FDR<0.05). (b) Selected chemokines and (c) inflammatory markers upregulated in plasma of AGMs following SARS-CoV-2 infection. (d) STRING protein-protein interaction network based on the 15 significantly differentially regulated plasma proteins at week 1. Proteins significantly regulated at p<0.05 with FDR >0.05 are highlighted in gray. Those with FDR<0.05 are highlighted in red. (e) Top functionally enriched networks associated with the 15 regulated plasma proteins in STRING. Proteins in biological processes associated with regulation of leukocyte migration are highlighted in green. Proteins with molecular functions associated with receptor ligand activities are highlighted in yellow. Proteins associated with chemokine receptor binding, and macrophage proliferation are highlighted in red. Proteins in KEGG pathways associated with regulation of leukocyte migration are highlighted in blue. Proteins that entered the network based on close associations are denoted by gray. (f) PANTHER 17.0 classification of differentially regulated proteins based on protein class, and (g) pathway.

We next examined whether these 15 differentially regulated analytes were interrelated or shared common pathways. We employed the STRING protein-protein analysis web-based tool using the default settings. Except for CDCP1 and SULT1A1 (ST1A1) there were high confidence interconnections between the regulated analytes, notably the chemokines CCL8, CCL19, CCL25, CXCL8, CX3CL1, and TNF and IL-18 (Fig. 3d). The top functional enriched networks revealed biological processes and molecular functions associated with regulation of leukocyte migration and receptor ligand activity. The top ranked local network clusters and KEGG pathways were related to chemokine receptors, macrophage proliferation, and viral protein interaction with cytokines and cytokine receptors (Fig. 3e, Table S2). We validated these networks using PANTHER v 17.0 revealing the protein class was dominated by intracellular signal molecules (Fig. 3f), with top pathways associated with inflammation mediated by chemokine and cytokine signaling, and apoptosis (Fig. 3g), supporting the STRING analysis. These data identify a specific dysregulation in inflammatory markers in the AGM metabolic-PASC model.

### 2.6 Glial cell line-derived neurotrophic factor (GDNF) remains persistently elevated following SARS-CoV-2 infection of AGM and associates with hyperglycemia

Increased serum concentrations of inflammatory cytokines correlate positively with fasting glucose levels in individuals with features of the metabolic syndrome^37^. We questioned whether early pathological inflammatory processes may be associated with early-onset, or persistence of hyperglycemia in our SARS-CoV-2 infected AGM long-COVID model. We conducted correlation analysis of blood glucose levels at week 1, 4, 8 and 16 against all analytes that were significantly (unadjusted p value <0.05) differentially regulated in the 10 unvaccinated animals at week 1 (Fig. 4a). We observed a modest but statistically significant correlation (p<0.05) between week 1 plasma CDCP1 levels and week 4 and 16 glucose levels, and a significant and strong positive correlation between week 1 plasma GDNF and week 4 glucose levels (Fig. 4a, Fig. S2). We explored the overall relationship between total plasma analytes and changes in plasma glucose concentrations over time in the unvaccinated group. We identified 8 analytes CCL25, GDNF, ADA, ST1A1, CXCL9, IL-10RB, FGF-19 and CDCP1 that positively and significantly correlated with plasma glucose across baseline, week 1, 4, and 12, and one analyte, IL-8, which had a significant negative association with glucose. CCL25 (r=0.57; p=0.0003; FDR=0.004), GDNF (r=0.55; p=0.0004; FDR=0.006) and ADA (r=0.44; p=0.007; FDR=0.049) maintained significance following BH correction (Table S3). The CCl25 and GDNF data are graphically represented (Fig. 4b). STRING functional protein interaction analysis of these 9 analytes identified CXCR chemokine receptor binding (FDR=0.019), chemokine receptor chemokines (FDR=0.009), and viral protein interaction with cytokines and cytokine receptors (FDR<0.0001) as top functional enrichments networks.

**Figure 4.**
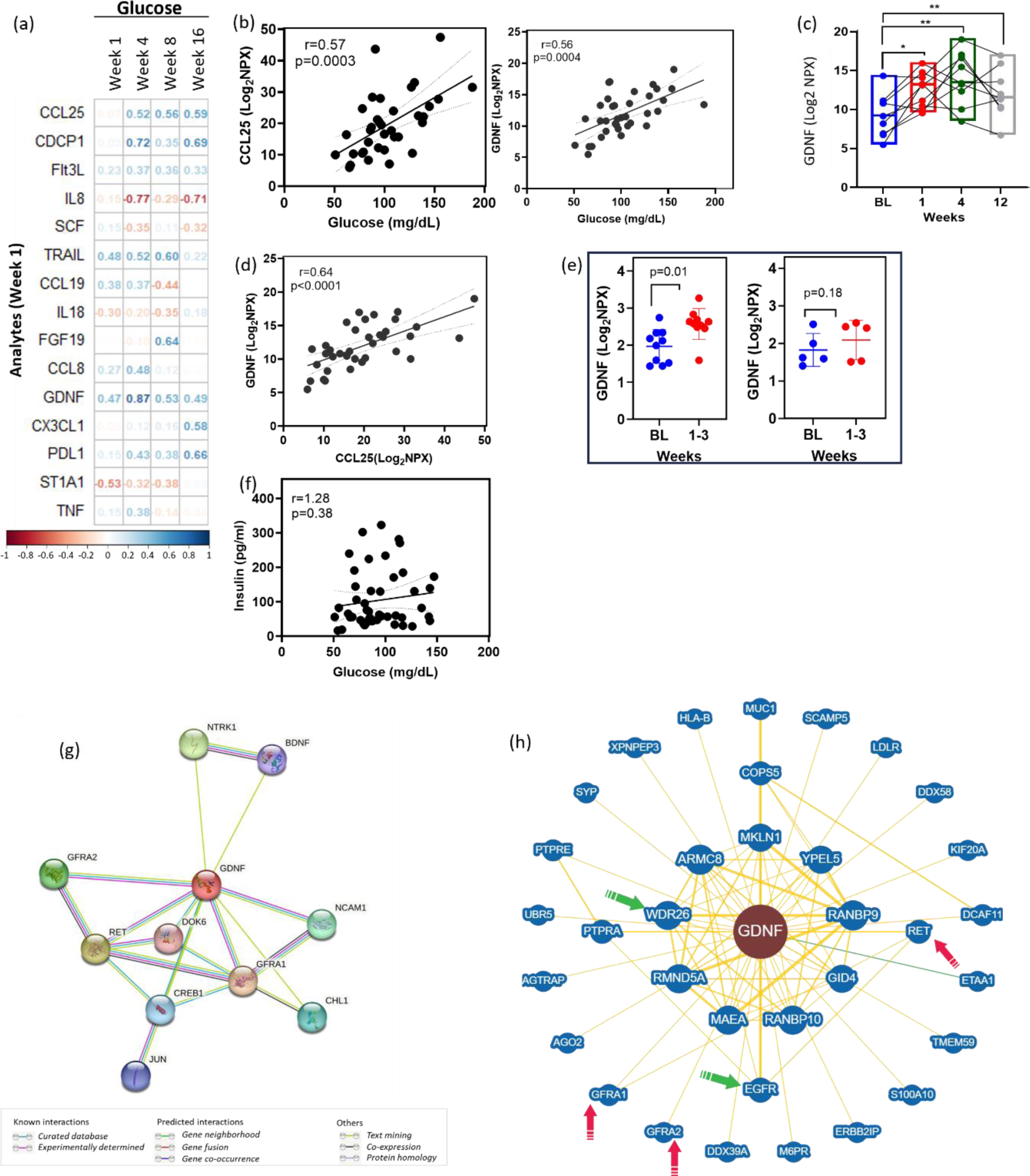
Plasma levels of GDNF and CCL25 correlate with blood glucose. (a) A correlation matrix depicting the r values between SARS-CoV-2-modulated analytes at week 1, and plasma glucose concentrations at various timepoints. Blue-colored correlations = positive correlations and red-colored correlations = negative correlations (raw p value correlation matrix in Fig. S2). Spearman’s rank correlation was used for statistical analysis. (b) Spearman’s correlations between plasma CCL19 or GDNF with glucose across baseline, week 1, 4 and 12 in the unvaccinated group. (c) Plasma GDNF in AGMs at various timepoints. Statistical significance was determined using the Wilcoxon signed rank test; * p<0.05, ** p<0.01. (d) Spearman’s correlations between plasma CCL19 and GDNF across baseline, week 1, 4 and 12 in the unvaccinated group. (e) Comparative analysis showing plasma GDNF levels in unvaccinated (left panel) and vaccinated (right panel) animals at baseline (BL) and weeks 1-3. (f) Spearman correlation between blood glucose and insulin across baseline, week 1, 4, 8, 16 and 12 in the unvaccinated group. (g) Unbiased STRING analysis, using custom setting with GDNF as the sole input. (h) BioGRID protein-protein analysis tool using customs settings and *Homo sapiens* database used to validate GDNF interactions (proteins of interest highlighted by arrows).

Of these analytes GDNF levels remained persistently and significantly higher above baseline up to week 12 p.i. (Fig. 4c), and correlated positively and significantly with plasma CCL25 (r=0.64, p<0.0001), a chemokine linked to T2D^29^ (Fig. 4d). We employed a confirmatory dataset comprising of plasma from both unvaccinated and vaccinated animals at baseline and 1-3 weeks and evaluated GDNF levels using the OLINK panel indicated above. We confirmed significantly increased plasma GDNF following SARS-CoV-2 infection of AGMs. At the same time points, there was no significant elevation in plasma GDNF in the vaccinated group (Fig. 4e). Interestingly, there was no significant correlation between blood insulin levels and glucose (Fig. 4f).

To gain more insights into the functionality of GDNF, we conducted unbiased analysis by using GDNF as the only search variable in STRING and set the interaction stringency to highest allowable confidence (0.90). Based on experimental evidence, databases, and text mining the analysis map revealed a high confidence interaction (0.999) between GDNF and its receptors GFRA1, and GFRA2, and the putative receptor RET receptor tyrosine kinase, which is involved in neuronal navigation and cell migration. There were also high confidence interactions between GDNF and neural cell adhesion molecule 1 (NCAM1; 0.936), a cell adhesion protein involved in neuron-neuron adhesion, and between GDNF and cyclic AMP-responsive element-binding protein 1 (CREB1), a transcription factor activated upon binding to the DNA cAMP response element (CRE) found in many viral promoters (Fig. 4g).

Finally, we used BioGRID, another protein-protein interaction database, to validate STRING’s GDNF interactions. The results confirmed our STRING analysis, showing high confidence interactions between GDNF and its receptors (red arrows), as well as with WDR26 (evidence: affinity capture-MS; green arrow; Fig. 4h), which serves as a scaffold to coordinate PI3K/AKT activation^38^, as well as regulating leucocyte migration^39^. GDNF also interacts strongly with epidermal growth factor receptor (EGFR; evidence: affinity capture-MS; green arrow) reported to regulate the severity of COVID-19 in patients^40^. Together, these results identify a potentially early and minimal host inflammatory signature dominated by elevation of plasma chemokines and the neurotropic factor GDNF, associated with impaired glucose homeostasis, in SARS-CoV-2 infected AGM.

### 2.7. Polyfunctional inflammatory CD4+ T cells positively correlate with blood glucose levels in SARS-CoV-2 infected AGMs

As shown, T cell hypersensitivity is increased following SARS-CoV-2 infection in AGMs. We therefore examined whether the degree of hypersensitive polyfunctional T cell responses correlate with blood glucose levels in unvaccinated AGMs. PBMCs were exposed to PMA/I for 6 hours and the levels of T cell activation and cytokine production were examined by flow cytometry. We imported data in FlowJo, and gated on singlets, live cells and CD45+CD3+ T cells (Fig. S3) and exported CSV - Scale values. We used the R package Spectre to identify unique polyfunctional/inflammatory populations (BL, week 1, 4, 8, 12, 18). In the unstimulated samples we defined two major populations, CD4 (cluster A) and CD8 (cluster B) shown by UMAPs (Fig. 5a-b, Fig. S4a). In the PMA/I treated PBMCs we defined nine populations (Fig. 5c), described in detail in Fig. S4b. The high expression (deep red) of activation and inflammatory markers confirmed activation status (Fig. 5d). There was a significant positive correlation between blood glucose levels and the level of the early activation marker CD69 on IL-2 producing memory CD4 T cells (Fig 5c, Population E; Fig. 5e), and a significant and positive correlation between blood glucose levels and the number of polyfunctional (TNF+ IFNγ+) naïve CD4 T cells (Fig. 5c, Population A; Fig. 5f). Plasma GDNF levels were also significantly correlated with the number of IL-17-producing activated (CD69+ IL-17+ IL-2+ TNF+) naïve CD4 T cells (Fig. 5c, Population C; Fig. 5g), as well as with the number of polyfunctional naïve CD4 T cells (Fig. 5h). These data identify populations of hypersensitive T cells induced ex-vivo by PMA/I, that correlates with hyperglycemia in SARS-CoV-2 infected AGMs.

**Figure 5.**
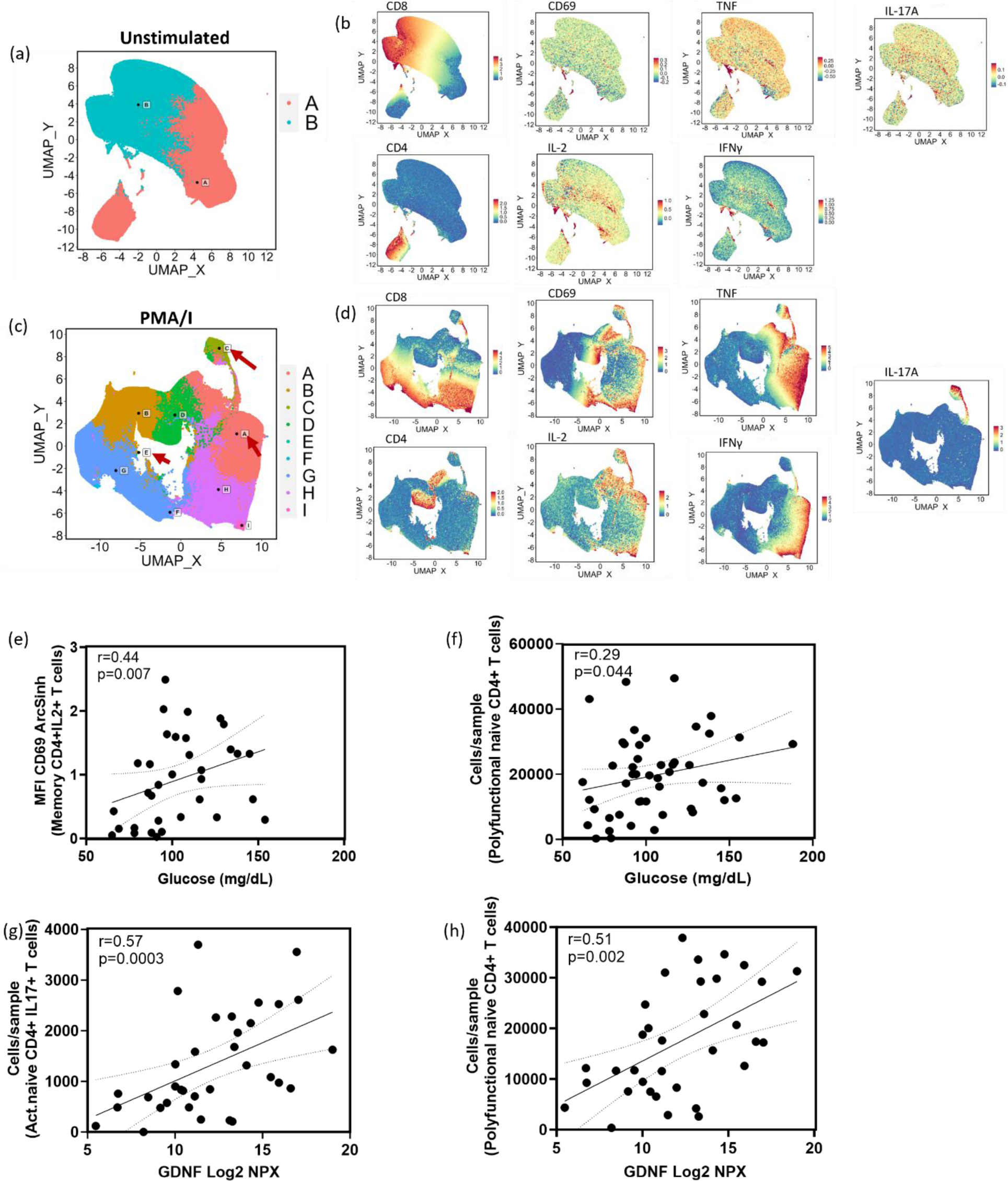
Multi-level analysis by Spectre identifies invitro-generated polyfunctional CD4 T cell populations correlating with plasma glucose and GDNF levels. (a) UMAP plot showing total CD4 (cluster A) and CD8 (cluster B) T cells in unstimulated PBMCs. (b) UMAPS showing expression of selected cell surface phenotypic, and inflammatory markers in unstimulated PBMCs. (c) UMAP plot showing clusters A-I (see cluster details in Fig. S4) within the CD4 and CD8 T cell populations of PMA/I treated PBMCs. (d) UMAPS showing expression of selected cell surface phenotypic, and inflammatory markers in PMA/I stimulated PBMCs. (e-f) Spearman correlations between blood glucose levels in unvaccinated animals and polyfunctional populations. (g-h) Spearman correlations between plasma GDNF levels in unvaccinated animals and polyfunctional populations.

### 2.8. Analysis of SARS-CoV-2 persistence in tissues 18 weeks post infection

Data suggest extrapulmonary presence of SARS-CoV-2 RNA in human tissues post-acute infection^41^, and replicating virus has been isolated from human hepatocytes from postmortem COVID-19 patients^20^. We therefore took advantage of RNAscope using an anti-sense probe targeting SARS-CoV-2 spike protein RNA (SARS-S) but observed no substantial percentage of cells containing spike RNA (SARS-S+ cells %) in duodenum (mean= 0.018%), liver (mean=0.005%), and pancreas (mean =0.012%) at 18 weeks p.i., as well as in historical sections (4 weeks p.i.) (mean: duodenum = 0.0037%, liver =0.0016%, pancreas =0.004%). Substantial SARS-S signals from lung samples of a SAR-CoV-2 infected AGM (4 weeks p.i.; Fig. 6j-k) support reliability of staining. For confirmatory study we conducted qPCR analysis on tissues collected 18 weeks p.i. We found no SARS-CoV-2 subgenomic N nor subgenomic E signals in the liver, duodenum, or pancreas, but low-levels of genomic N signals (Ct mean 30.4 and 31.6) in the duodenum of two of the 15 animals (data not shown). We conducted SARS-CoV-2 immunohistochemistry on duodenum collected 18 weeks p.i. but found no positive signals (Fig 6. l-m). Together these data suggest no significant long-term persistence of replicating virus in the liver, pancreas or duodenum in our AGM PASC model.

**Figure 6.**
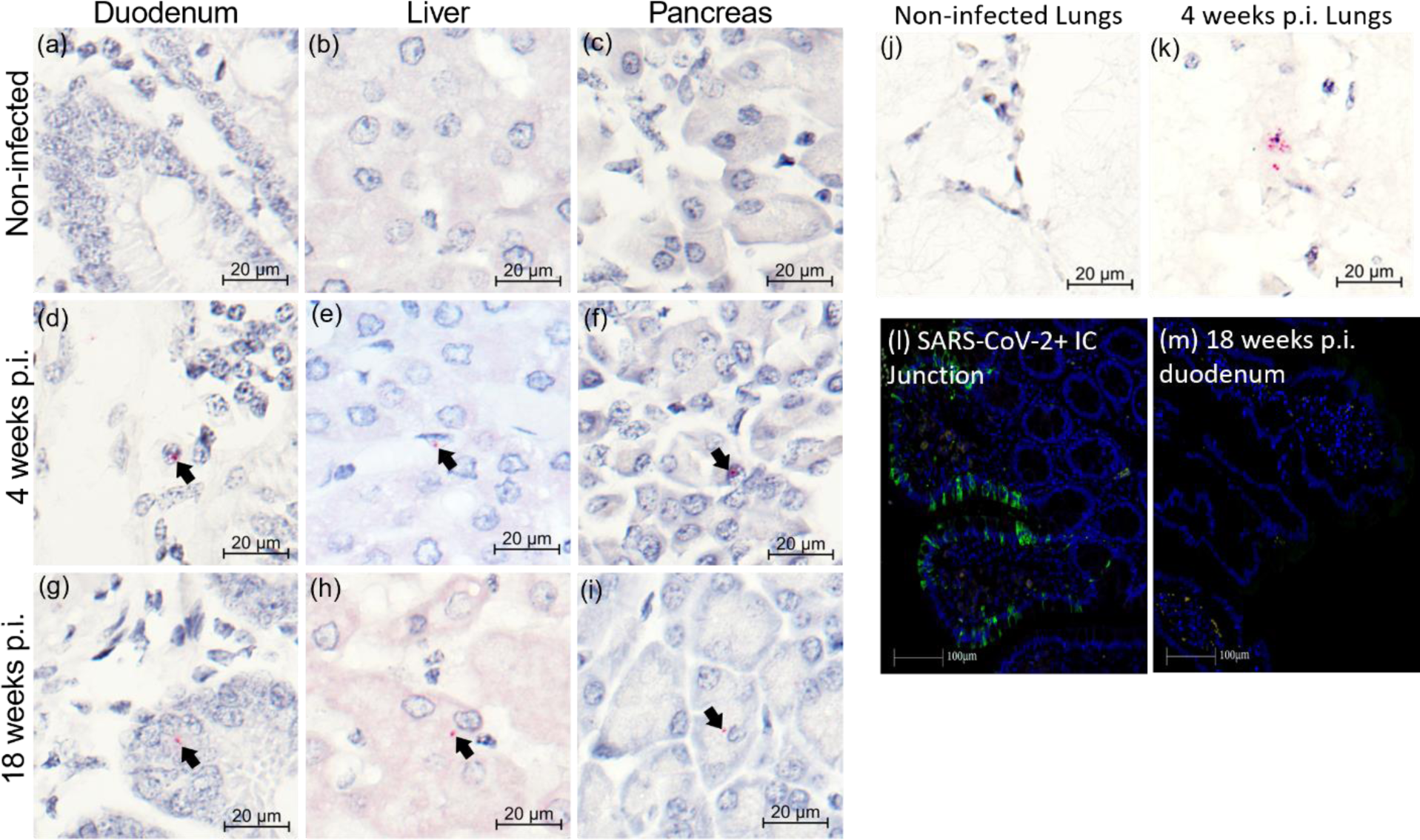
Persistence of SARS-CoV-2 RNA in tissues of AGMs up to 18 weeks post infection. (a-c) Representative images of RNAscope RED using the SARS-Spike (S) probe in duodenum, liver, and pancreas in non-infected animals, (d-f) 4 weeks post infection (p.i.), and (g-i) 18 weeks p.i. RNAscope RED was used to visualize the SARS-S expression frequency in the tissues, counterstained with hematoxylin. Black arrows note the presence of SARS-S copies as red dots by colorimetric RNAscope. (j-k) Representative validation images of RNAscope RED using the SARS-S probe in lungs of non-infected, and 4 weeks p.i. animals. Tissues were counterstained with hematoxylin. (l-m) Fluorescent immunohistochemistry in a SARS-CoV-2 positive control (l) and a representative animal 18 weeks p.i. (m). Blue = DAPI; Green = anti-SARS antibody.

### 2.9. Absence of severe lung inflammation and injury during long-term follow-up in SARS-CoV-2 infected AGM

We previously reported two of four AGM (2/4) exposed to SARS-CoV-2 isolate USA-WA1/2020 progressed to acute respiratory distress syndrome (ARDS) by day 8 and 22 p.i. and exhibited diffuse alveolar damage and bronchointerstitial pneumonia^25^. We therefore examined whether viral-induced high grade lung inflammation is present at our 18 weeks study endpoint. We conducted histopathological analysis of lungs from all study animals at necropsy (n=15, end point 18 weeks p.i.), and included 3 uninfected animals, and four infected from a previous study with endpoint between 3 and 4 weeks p.i. The inflammation grades were scored 1 to 4 (minimal to severe) based on indices of inflammation, utilizing the numbers of inflammatory cells, the degree of fibrous connective tissue formation (Fig. S5a, Panels A-D), and the presence of pneumocyte type II hyperplasia (Fig. S5b, Panels A-B). Analysis of the right lower lung shows minimal inflammation in all except one animal in the long-COVID study group. Two of the four animals from the short-term study group had severe inflammation. We generated a composite inflammation score by pooling scores from the right anterior upper, right middle dorsal and right lower lung but found no overall severe inflammation above that of the three uninfected AGM controls (Fig. S5c). There was no evidence of aspirated pneumonia or euthanasia artifacts in the two AGMs from the short-term study, with severe lung inflammation, suggesting this may be SARS-CoV-2-related. The data suggest that at least in the animals examined by our inflammation measurements there was no general severe lung inflammation at 18 weeks p.i.

### 2.10. Absence of severe pancreatic inflammation and injury during long-term follow-up in SARS-CoV-2 infected AGM

Extrapulmonary manifestations of SARS-CoV-2 infection includes infection of the human exocrine and endocrine pancreas, causing morphological changes that may contribute to impaired glucose homoeostasis^26, 27, 34^. We examined the inflammation and fibrosis status of hematoxylin and eosin (H&E) stained pancreatic cross sections, obtained at necropsy, and found none to minimal inflammation, like the two uninfected historic samples. There was no evidence of pancreatic fibrosis in 13 animals, while 2 demonstrated mild fibrosis. Representative images are shown (Fig. S6). Taken together, we saw no evidence to support significant long-term morphological defects of the pancreas.

### 2.11 SARS-CoV-2 infection is associated with increased liver glycogen levels

Although previous reports show SAR-CoV-2 infection of pancreatic β-cells in humans and AGMs, infection in humans may be independently associated with hyperglycemia regardless of β-cell function^20^. SARS-CoV-2 infected hepatocytes have raised glucose production through increased gluconeogenesis, a potential cause of hyperglycemia in infected humans^20^. Regardless of the underlying source, elevated blood glucose and hepatic glucose uptake may provide substrate for hepatic glycogen synthesis^42^ (Fig. 7a). We therefore quantified liver glycogen in uninfected and 4-weeks p.i. SARS-CoV-2 (short term infection) infected historical samples, as well as at necropsy (long-term infection) using Periodic acid–Schiff (PAS) staining. Representative staining is shown without diastase (Fig. 8b, upper panel), and with diastase (Fig. 8b, lower panel) to confirm staining specificity. A trend for increased liver glycogen in infected animals was observed (Fig. 7c), reaching significance in the longer-term infected unvaccinated group. Liver glycogen levels correlated positively with blood glucose levels at week 8 (Fig. 7d) and 12 (Fig. 7e-f) p.i. Of note, we saw no evidence of hepatic steatosis or fibrosis at necropsy.

**Figure 7.**
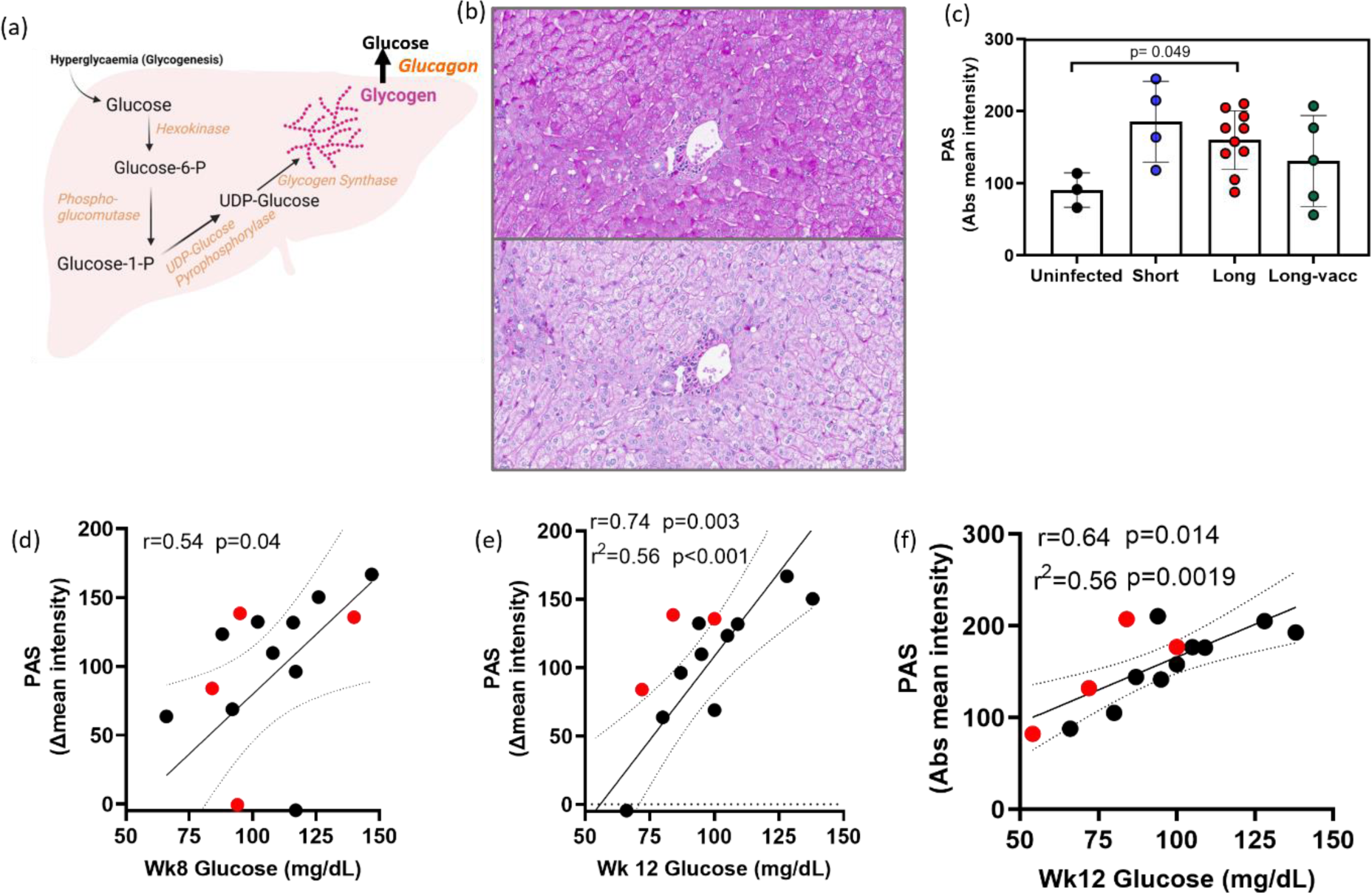
Liver glycogen levels and relationship with blood glucose. (a) A model of liver glucose homeostasis during SARS-CoV-2 infection. Elevated blood glucose triggers glycogenolysis, which stores glucose into glycogen. Increased glucagon levels in SAR-CoV-2 infection stimulates the conversion of stored liver glycogen back to glucose perpetuating a vicious cycle. (b) Representative image of Periodic acid–Schiff staining of AGM liver at 18 weeks post infection (p.i.) without diastase (upper panel), and with diastase (lower panel). (c) Glycogen levels in livers from uninfected AGMs, 4-week p.i (short term) and at 18-week p.i (long term unvaccinated or vaccinated). (d-f) Spearman’s correlation between blood glucose at various weeks and hepatic glycogen at 18 weeks p.i. in unvaccinated (black dots) or vaccinated (red dots). Intensity of stain was quantified as either delta mean intensity (difference between diastase treatment and untreated), or absolute intensity.

## 3. Discussion

Collateral damage by early host antiviral responses against SARS-CoV-2 may underlie the severity and clinical presentation of acute COVID-19 symptoms. Pre-existing or virus-induced metabolic provocations may promote or exacerbate these acute symptoms, as well as contribute to the development and long-term phenotype of metabolic syndromes such as MAFLD^13^, and hyperglycemia^35^. Here we describe the first study to systemically evaluate, in non-human primates (NHPs), the temporal changes of hyperglycemia (metabolic PASC) over an 18-week period. We observed, in SARS-CoV-2 infected AGMs, a multitude of immunologic and metabolic changes that reflect those previously reported in humans in the acute and post-acute phase of COVID-19.

Our data demonstrate that SARS-CoV-2 infection of AGMs is associated with early onset hyperglycemia, which persisted for at least 18 weeks p.i. We have previously reported that early immune changes contribute to adverse pathological events in SARS-CoV-2 infected NHPs^43^. Here, we identified a set of plasma analytes, many differentially increased at week 1 p.i., that correlated positively and significantly with plasma glucose levels over time. The signature of these regulated analytes was defined by a preponderance of chemokines and several inflammatory-related proteins. Using protein-protein interaction tools and gene-ontology analyses we have discovered that the top functionally enriched networks associated with these analytes are related to leukocyte migration, chemotaxis, macrophage proliferation and viral protein interaction with cytokines. Amongst these analytes, we noted CCL25 and GDNF to be significantly elevated at week 1 p.i. and beyond and correlated positively with blood glucose levels across all timepoints evaluated. Thus, using the highly specific PEA technology we identified a set of differentially regulated plasma proteins associated with early and persistent impairment of glucose homeostasis in SARS-CoV-2 infected AGMs. Additionally, we report increased T cell hypersensitivity during infection, with polyfunctional responses to ex-vivo PMA/I stimulation positively correlating with glucose levels. Finally, while the magnitude of antibody responses was similar in the vaccinated and unvaccinated groups, we show that the vaccination of five animals 4 days p.i. was associated with a consistent and significantly lower blood glucose level over the study period.

Previous reports show that elevated glucose levels favor SARS-CoV-2 infection and monocyte pro-inflammatory responses^44^, as well as elevating the risks of severe COVID-19 progression and increased fatality^17, 18^. In addition, T cell hypersensitivity towards polyclonal stimulation (PMA/I) is seen in severe and extreme cases of COVID-19^12^. Hyperglycemia may be caused by insulin resistance, dysfunctional pancreatic β-cells, impaired glucose clearance or increased glucose disposal by the liver through gluconeogenesis or glycogenolysis^20^. We have previously shown in a short-term 4-week study, in two AGMs with demonstrable SARS-CoV-2 infection of the pancreatic ductal and endothelial cells, that infection was associated with pancreatic thrombofibrosis and new-onset diabetes^34^. The focus of this AGM PASC study precludes dissecting the specific mechanisms by which glucose homeostasis is regulated during infection. Nonetheless, pathological elevation of blood glucose in moderate and severe COVID-19 patients has been linked to enhanced hepatic gluconeogenesis by the activity of the Golgi protein GP73, which is found to be elevated in the plasma of infected patients^35^. Although it remains controversial whether SARS-CoV-2 can replicate in hepatocytes, data show that these cells express the chaperone glucose-regulated protein 78 (GRP78), a putative SARS-CoV-2 entry factor, as well as low levels of ACE2 protein, which co-localized with spike protein. Moreover, the co-localization of spike protein with viral RNA, and ex-vivo infection assays support the inference that SARS-CoV-2 can replicate in the liver^20^. Although the infection appears non-cytopathic, it was proposed that SARS-CoV-2, including the major gamma, delta, and omicron variants, can stimulate hepatic glucose production and disposal through increased activity of the rate limiting gluconeogenic enzyme phosphoenolpyruvate carboxykinase (PEPCK)^20^. The key focus of this work was to ascertain whether AGMs infected with SARS-CoV-2 are feasible models to study metabolic PACS, so detailed mechanistic analyses are beyond this study. However, we observed elevated glycogen levels in hepatocytes of infected AGMs at necropsy, which positively correlated with blood glucose levels. On the contrary we saw no substantial amount of SARS-CoV-2 nucleic acid or proteins in the liver or pancreas 18 weeks p.i. Glucagon, which can promote the breakdown of stored hepatic glycogen to glucose, is increased in COVID-19 patients’ plasma^20^, but we have not interrogated this connection in our model. Nonetheless this supports the idea that non-insulin, and non-pancreatic related mechanisms may partially regulate glucometabolic control during SARS-CoV-2 infection ^35^.

To gain insights into other potential processes driving COVID-19 related hyperglycemia, we assessed a panel of blood inflammatory/metabolic analytes at baseline and at set study time points. The signature of proteins differentially increased at week 1, and associated with plasma glucose levels over time, was biased towards those involved in leukocyte migration and chemokine receptor binding. Some of these include CCL8, CCL19, and the gut homing chemokine CCL25 that correlated strongly and positively with blood glucose levels across multiple weeks.

Recent studies show plasma chemokines to be critical factors that control COVID-19 severity^45, 46^. Accordingly, CCL25 has been found to be elevated in the plasma of COVID-19 patients^47^, and a large-scale genome wide association study, involving patients with severe COVID-19, identified mutations in several chemokine receptors, including the CCL25 receptor CCR9 as a major risk factor for developing severe COVID-19^48^. We found no evidence of long-term pancreatic damage, however the elevation of CCL25 could potentially impair insulin secretion from the pancreas.

Intriguingly, while anti-SARS-CoV-2 antibodies may counter viral replication, antibodies against inflammatory mediators such as CCL25, may be more effective at counteracting development of both acute and long-COVID symptoms. We found our set of differentially regulated chemokines (CCL8, CCL19 and CCL25) intriguing because autoantibodies against CCL8 are augmented in long-term convalescent COVID-19 individuals, and elevated antibodies against the COVID-19 signature chemokine CCL19^46^ are documented with high confidence in both acute and long-term COVID-19 phases compared to uninfected controls^49^. Interestingly, autoantibodies against CCL25 are augmented in mild COVID-19 patients compared to those requiring hospitalization^49^. Convalescents exhibiting PASC at 12 months have a significantly lower cumulative level of anti-chemokine antibodies at six months compared to those who reported no PASC^49^. In our STRING protein-protein analysis of significantly regulated analytes at week 1, we found strong interactions between CCL8, CCL19, CCL25, TNF and IL-18 confirming a link between this chemokine signature and inflammatory responses in our infected AGM model, likely contributing to early adverse and long-term pathological events.

Some evidence suggests a link between inflammatory processes and COVID-19 related metabolic diseases, but our understanding of the underlying mechanisms remains limited. Chemokines are best known for their role in immune cell trafficking to sites of infection and as mediators of inflammation and tissue repair. However, recent reports have linked their activity to features of metabolic syndrome such as insulin resistance and T2D. CCL25 acting via its receptor CCR9 impairs β-cell function and inhibits glucose-induced insulin secretion^29^. Moreover, CCR9 has been implicated in the pathogenesis of T2D by modulating small intestine permeability and inflammation^50^. Although we found a strong and significant positive relationship between blood glucose and CCL25 levels, linking this causatively to glucose homeostasis in COVID-19 requires further investigation.

Besides the classic chemokines, we discovered GDNF to positively associate with plasma glucose levels over time. GDNF is a neurotrophic factor belonging to the transforming growth factor-β (TGF-β) superfamily, which plays a key role in the nervous system, and the pathogenesis of mood disorders. GDNF is a known canonical RET ligand, validated by our STRING analysis, which demonstrates a high confidence interaction between GDNF and RET. While the role for GDNF in peripheral glucose metabolism is unclear, another RET ligand GDF15, also belonging to the TGF-β superfamily, is known to regulate systemic metabolic homeostasis, and is a correlative biomarker for metabolic syndrome^51, 52^. Moreover, GDF15 binds with high affinity to GDNF family receptor α-like (GFRAL), an interaction required for GDF15-RET binding^53, 54^ and may represent a compensatory checkpoint during conditions of high metabolic stress such as SARS-CoV-2 infection. In fact, GDF15 levels are elevated in SARS-CoV-2-infected patients and are significantly associated with worse clinical outcomes^55^.

GDNF has been shown to reverse the pathological effects of hyperglycemia on enteric neuronal survival via activation of the PI3K-Akt pathway^56^, a signaling cascade that regulates Glut1 and Glut4 mediated glucose uptake into cells^57, 58^. Recently, nutritional regulation of GDNF has been suggested, in which its expression was enhanced by glucose^59^. Interestingly, in protein-protein interaction analysis, besides its receptors, GDNF also interacts with Neural Cell Adhesion Molecule 1 (NCAM1) with high confidence, confirming its potential role as a chemotaxis factor, especially for epithelial and enteric neural cells essential for maintaining gut wall integrity^60, 61^. Since GDNF levels correlate inversely with plasma glucose in T2D patients^62^, and have been shown to improve glucose tolerance and increase β-cell mass in vitro and in vivo^63^. The increased GDNF in our studies is likely an adaptive response. The precise role of GDNF in COVID-19 related pathologies is unknown but may represent a compensatory immunometabolic adaptation related to changes in energy metabolism in infected AGMs. In conclusion, we show SARS-CoV-2 infected AGMs exhibit many virologic, immunologic and metabolic features observed in infected humans and may represent a useful model to interrogate early, and persistent factors associated with metabolic PASC. We identify GDNF and several plasma analytes, dominated by chemokines, that are associated with hyperglycemia over several months p.i. We provide leads involving inflammatory processes, as well as potential dysregulated liver glucose homeostasis, that warrants further investigation to improve our understanding of how early inflammatory and metabolic responses against SARS-CoV-2 infection influence its severity and long-term metabolic complications. Such understanding may provide the basis for exploring autoantibodies of chemokines/metabolic-regulating factors to treat and prevent long-COVID. Since mRNA vaccines may elicit an immune response within hours and induce humoral immunity within 5 days of administration^64^, it is plausible that such responses may offer favorable immunometabolic benefits prior to multiorgan distribution of SARS-CoV-2 produced by viral shedding from the lungs into body fluids. Intriguingly, in an observational cohort study of 15 million people COVID-19 vaccination reduced the incidence of long-term diabetes significantly^65^. Our observation of better glycemic control in the vaccinated group requires further studies to evaluate the potential benefits of vaccination during the acute phase of infection^66^.

## 4. Methods

### 4.1 Study approval

This study was reviewed and approved by the institutional Animal Care and Use Committee of Tulane University. Animals were cared for in accordance with the NIH’s *Guide for the Care and Use of Laboratory Animals*. Procedures for handling and BSL2, and BSL3 containment of animals were approved by the Tulane University Institutional Biosafety Committee. The Tulane National Primate Research Center is fully accredited by the Association for Assessment and Accreditation of Laboratory Animal Care.

### 4.2 Animals and infection procedure

Procedures are in accordance with those we have previously reported^25^. Briefly, we exposed 15 African green monkeys (*Chlorocebus aethiops sabaeus*; 13 females, 2 males) aged 7.92 to 19.32 years, to SARS-CoV-2 strain 2019-nCoV/USA-WA1/2020 at ∼1e6 TCID50 via intranasal (0.5mL/nares), and intratracheal (1mL) routes. Except for one (PB24), obtained from the NIH via the Wake Forest breeding colony, all animals were of Caribbean origin (wild-caught) purchased from Bioqual (MD, USA). Ten animals (9 females) were studied during the natural course of SARS-CoV-2 infection and 5 animals (4 females) received the BNT162b2 Pfizer/BioNTech vaccine 4-days post infection. Animals were monitored daily for 18 weeks. Animals were anesthetized with telazol tiletamine hydrochloride and zolazepam hydrochloride (5 to 8 mg/kg intramuscular; Tiletamine–zolazepam, Zoetis, Kalamazoo, MI) and buprenorphine hydrochloride (0.03 mg/kg).

### 4.3 Blood chemistry and hematological analysis

A comprehensive biochemistry analysis on blood EDTA-collected serum was performed at the TNPRC clinical lab, using the Beckman Coulter AU480, according to the manufacturer’s instructions. The panel included albumin, glucose, cholesterol, triglycerides, aspartate aminotransferase (AST), alanine aminotransferase, (ALT), blood urea nitrogen (BUN), alkaline phosphatase (ALP), and lactate dehydrogenase (LDH). Hematological analysis on whole blood, including absolute quantification, and percentages of neutrophils, monocytes, lymphocytes, and eosinophils were performed on the Sysmex NX-V-1000 Hematology Analyzer.

### 4.4 Virological analysis: Genomic and subgenomic RNA quantitation

Pre- and post-exposure samples of mucosal swabs (nasal and pharyngeal brush) were obtained for virological analysis. For RNA extraction from swab samples 200µL of 1× DNA/RNA Shield (Zymo) was added to each swab and RNA was extracted using the Zymo Quick RNA Viral Kit (Zymo) according to manufacturer’s instructions. Samples were eluted in 50µL volume. Subgenomic and genomic SARS-CoV-2 mRNA were quantified as previously described using appropriate primers/probes, and cycling conditions ^67, 68^, with the exception that, the probe for detection of the subgenomic nucleocapsid RNA was modified as described^69^. Briefly, qPCR analysis was conducted on a QuantStudio 6 (Thermo Scientific, USA) using TaqPath master mix (Thermo Scientific, USA). Signals were compared to a standard curve generated using in vitro transcribed RNA of each sequence diluted from 10^8^ down to 10 copies. Positive controls consisted of SARS-CoV-2 infected VeroE6 cell lysate. Viral copies per swab were calculated by multiplying mean copies per well by volume of swab extract.

### 4.5 Vaccination

10 animals were studied during the natural course of SARS-CoV-2 infection and 5 animals receive one dose of the BNT162b2 (Pfizer/BioNTech) vaccine 4 days post infection.

### 4.6 Antibody response analysis

Detection of SARS-CoV-2 spike (S), spike S1 RBD (S1 RBD), and nucleocapsid (N) proteins were performed using MSD S-PLEX CoV-2, MSD S-PLEX CoV-2 S1 RBD and MSD S-PLEX CoV-2 N assay kits for IgA, IgM, and IgG antibodies (Meso Scale Discovery, Rockville, MD). The assays were done according to the manufacturer’s instructions. Plasma samples were diluted 1/500-fold in the assay buffer provided. The plates were read using the MESO QuickPlex SQ 120MM reader. Sample quantitation was achieved using a calibration curve generated using a recombinant antigen standard. During analysis, any concentrations below the limit of detection (LOD) were assigned the LOD value, and any concentrations above the highest calibration standard were assigned its value.

### 4.7 Hypersensitive T cell response

Cell Stimulation Cocktail (eBioscience) containing phorbol 12-myristate 13-acetate (PMA), ionomycin, brefeldin A and monensin were used ×1 to stimulate PBMCs. Briefly, PBMCs were thawed, stimulated, and incubated for 6 hours in supplemented RPMI-1640 medium [10% human serum, penicillin/streptomycin (Invitrogen), 2 mmol/l L-glutamine (Invitrogen, Carlsbad, California, USA)] at 37°C, 5% CO_2_. PBMCs were stained with Zombie Aqua (Biolegend, San Diego, CA) for live/dead cell gating, and surface stained using the following pre-titrated antibodies from BD Biosciences (San Jose, California, USA) or BioLegend: CD45-PerCP (BD, clone D058-1283), CD3-BV650 (BD, clone SP34-2), CD4-BV786 (BD, clone L200), CD8-BUV737 (BD, clone RPA-T8), CD28-BV605 (clone CD28.2), CD95-BV711 (clone DX2), CD69-PE-CF594 (clone FN50) and CD107a-BUV395 (BD, clone H4A3). Except for CD45-PerCP (NHP), all antibodies have verified reactivity to humans, and cross reactivity with NHPs. Cells were fixed and permeabilized using BD Fixation/Permeabilization solution (BD Biosciences) and stained for intracellular cytokines using the following antibodies from BioLegend: IL-2-BB700 (clone MQ1-17H12), TNF-APC (clone MAb11), IFNy-PE-Cy7 (clone 4S.B3), IL-4-BV421 (clone 8D4-8), IL-17A-PE (clone BL168). Cells were fixed in 2% PFA and acquired on a BD FACSymphony™ by the Flow cytometry core (TNPRC). Data were analyzed using FlowJo software, version 10.8.1 (Tree Star Inc., Ashland, Oregon, USA) or used for Spectre analysis in R (v4.2.1).^70, 71^

### 4.8 Plasma analyte (OLINK) analysis

Plasma analytes were analyzed using a proximity extension assay (Olink, Proteomics)^72^. Plasma was collected from freshly collected blood in EDTA anticoagulant tubes and centrifuged at 650 × g for 10 minutes. The plasma was aliquoted to minimize freeze-thawing and stored at −80°C. Samples were processed at the OLINK Analysis Services Lab in Waltham (MA, USA) or within the High Containment Research Performance Core (TNPRC). The Olink® Target 96 inflammation panel (Olink Proteomics AB, Uppsala, Sweden) was used to measure proteins following manufacturer’s instructions. In brief, pairs of oligonucleotide-labeled antibody probes are mixed with plasma to allow binding to their targeted protein. Oligonucleotides will hybridize in a pair-wise manner if the two probes are brought in proximity. Reaction mixture containing DNA polymerase allows proximity-dependent DNA polymerization and creating a unique PCR target sequence. The amplified DNA sequence is quantified, quality controlled and normalized using internal control and calibrators. Protein levels are expressed as arbitrary units NPX values.

### 4.9 Insulin determination

Insulin plasma levels were measured using the monkey insulin ELISA kit (AssayGenie, Dublin Ireland). The essay was conducted according to manufacturer’s instructions using 1/8 diluted samples.

### 4.10 RNA Scope Analysis

Formalin-fixed paraffin embedded (FFPE) tissues were collected at necropsy and sectioned at 5 μm. In-situ hybridization (ISH) was conducted using RNAscope® 2.5 High Definition (HD) RED Assay Kit (Advanced Cell Diagnostics), according to the manufacturer’s directions. Briefly, FFPE tissue sections were deparaffinized in xylenes and dried, followed by incubation with hydrogen peroxide. Heat-mediated antigen retrieval was carried out in a steamer with the provided kit buffer. Samples were treated with the kit-provided protease and hybridized with the V-nCoV2019-S probe in a HybEZ oven (Advanced Cell Diagnostics). All washes were performed with the kit wash buffer. Signal amplification was accomplished with six successive AMP solutions and the kit-provided Fast Red dye. Slides were counterstained with hematoxylin. Control slides were included in every run to confirm specificity of staining and assess background.

For imaging and quantitation, brightfield images were acquired using the Axio Observer 7 (Zeiss), equipped with ZEN blue edition software (v3.6.096.08000). Images were subjected to brightness and contrast enhancement in Photoshop (Adobe, v24.4.0) applied to the entire image. Slides were scanned with the Axio Scan.Z1 digital slide scanner (Zeiss). Scanned files were analyzed with HALO (Indica Labs, v3.4.2986.151) algorithm ISH (v4.1.3) for a non-biased measurement of copies on a cell-by-cell basis. The ISH algorithm was run in annotations specific to the tissue section of interest, using hematoxylin-stained nuclei to quantify the number of cells and Fast Red intensity and size accounted for positivity of the probe within the cell. Each resulting count was assessed individually, and all false positives were excluded.

### 4.11 SARS-CoV-2 immunohistochemistry

Formalin-fixed, paraffin-embedded tissue sections were deparaffined using standard procedures followed by heat (microwave) induced antigen retrieval in a high pH solution (Vector Labs H-3301), rinsed in hot water and placed in heated low pH solution (Vector Labs H-3300) and allowed to cool to room temperature. Sections were washed in phosphate buffered saline, blocked with 10% normal goat serum (NGS) for 40 minutes and incubated for 60 minutes with a 1:1000 dilution of guinea pig anti-SARS antibody (BEI NR-10361). The slides were incubated with a 1:1000 diluted goat anti-guinea pig secondary Alexa Fluor 488 conjugated antibody (Invitrogen A11073) for 40 minutes. Nuclei were labelled with DAPI (4’,6-diamidino-2-phenylindole). Images were taken with a Zeiss Axio.Z1 Slide Scanner and analyzed using HALO HighPlex FL v4.1.3 (Indica Labs).

### 4.12 Periodic Acid Schiff

Slides were deparaffinized on an auto Stainer (Histology Core, TNRPC) and subjected to Periodic Acid Schiff Hematoxylin stain (PASH) for glycogen. Briefly, slides were either treated with a 1% solution of diastase (control) or not, and samples oxidized with 1% Periodic Acid (Poly Scientific) for 10 minutes, washed, placed in Schiff’s reagent for 10 minutes, counterstained with hematoxylin and eosin (H&E), and cover slipped using standard procedures. Stain intensities were quantified using automated settings in ImageJ1.53t (Fiji)^73^.

### 4.13 Histopathological scoring

All relevant tissue samples were fixed, paraffin-embedded and stained with H&E for histopathological analysis of inflammation, fibrosis, or other relevant pathologies by an experienced, board-certified pathologist.

### 4.14 Bioinformatics (Protein-Protein interactions & pathway analysis)

Protein-protein interaction analysis was performed mainly using STRING (Search Tool for the Retrieval of Interacting Genes/Proteins (v 11.5)^74^. The *Homo sapiens* database was selected for the input data search. Unless otherwise noted, the default settings were used, including 10 as the maximum number of interactors to show, and a medium confidence of 0.400. The top 3 functional enriched networks with FDR <0.05 are reported. Functional classification of proteins was determined using PANTHER (Protein Analysis Through Evolutionary Relationships; v 17.0; http://www.pantherdb.org/) classification system ^75, 76^. BioGRID (v 4.4; https://thebiogrid.org/), a biomedical interaction repository, was also used to validate interactions between entries.

### 4.15 Spectre analysis

Spectre analysis was conducted as previously published^70^. The “CSV - Scale values” for each cell population of interest were exported from FlowJo. The exported data were analyzed using the Spectre R package workflow. Briefly, the flow cytometry standard (FCS) data were arcsinh transformed and clustered with FlowSOM. The clustered data were down sampled, and dimensionality reduction was performed with UMAP. The clusters were manually labelled as desired. Finally, the clusters were used to generate summary statistics, which were used for further statistical analysis.

### 4.16 Statistical analysis and packages

R was used to perform statistical analysis, principal components analysis, and ggplot2 (v 3.3.3) was used to create principal component plot, heat maps, and correlation matrices. To visualize the longitudinal changes in antibody levels over time, the data were log10-transformed. We employed the LOESS (Locally Estimated Scatterplot Smoothing) regression method of the ggplot2 package for its ability to create smooth curves that effectively capture underlying trends and variations in the data. By utilizing ggplot2 in R, we enhanced scatterplots with these smooth LOESS regression curves, providing a clear representation of the evolving trends in both variables over time.

Additional graphs were created using GraphPad Prism (v 9.0). Wilcoxon matched pairs signed-rank test was used for paired analysis, Spearman’s rank correlation tests for correlation, and PERMANOVA (in R; package vegan) for non-parametric ANOVA with permutations. Statistical significance is indicated by p< 0.05.

## Acknowledgements

This study was supported by the NIH-funded base grant to the TNPRC (P51OD011104), and NIH supplement (3P51OD011104-61S1). Additional support was provided by TUHS Auxiliary Endowment for Excellence at TNRPC to C.S.P. NIH S10 OD026800, which was awarded to support the TNRPC Flow Cytometry Core Facilities.

## Contributions

J.R., J.D., R.B., T.F., R.V.B., A.B. and K.R.L. designed animal study. C.S.P. and J. R. designed laboratory study. C.S.P., C.C., N.G., G.L., N. J. M., P. D., K.M.G., L. H., K. W., M.F., K. M. G. and K.B. participated in tissue acquisition and processing and performed experiments. C.S.P., C.P., A.S. and T.F., M A-M., R. T., and P. D. analyzed data. J. C. M. provided significant and substantive data analysis support. C.S.P., J.R., C.P., A.S., R.T., and T.F. interpreted data. C.S.P., J.R., C.P., M. A-M., A.S., C.M., and T.F. provided significant intellectual input. C.S.P. wrote the manuscript. J.R., C.P. and M. A-M. provided critical, and substantive intellectual editing. C.S.P., C.P., T.F., A.S., and M. A-M. prepared manuscript figures. C.K provides quality assurance and data management tasks. P.D., N. J. M., C. C., R.T., and T.F edited manuscript. All authors approved the manuscript.

## Supplementary figures

**Figure S1.**
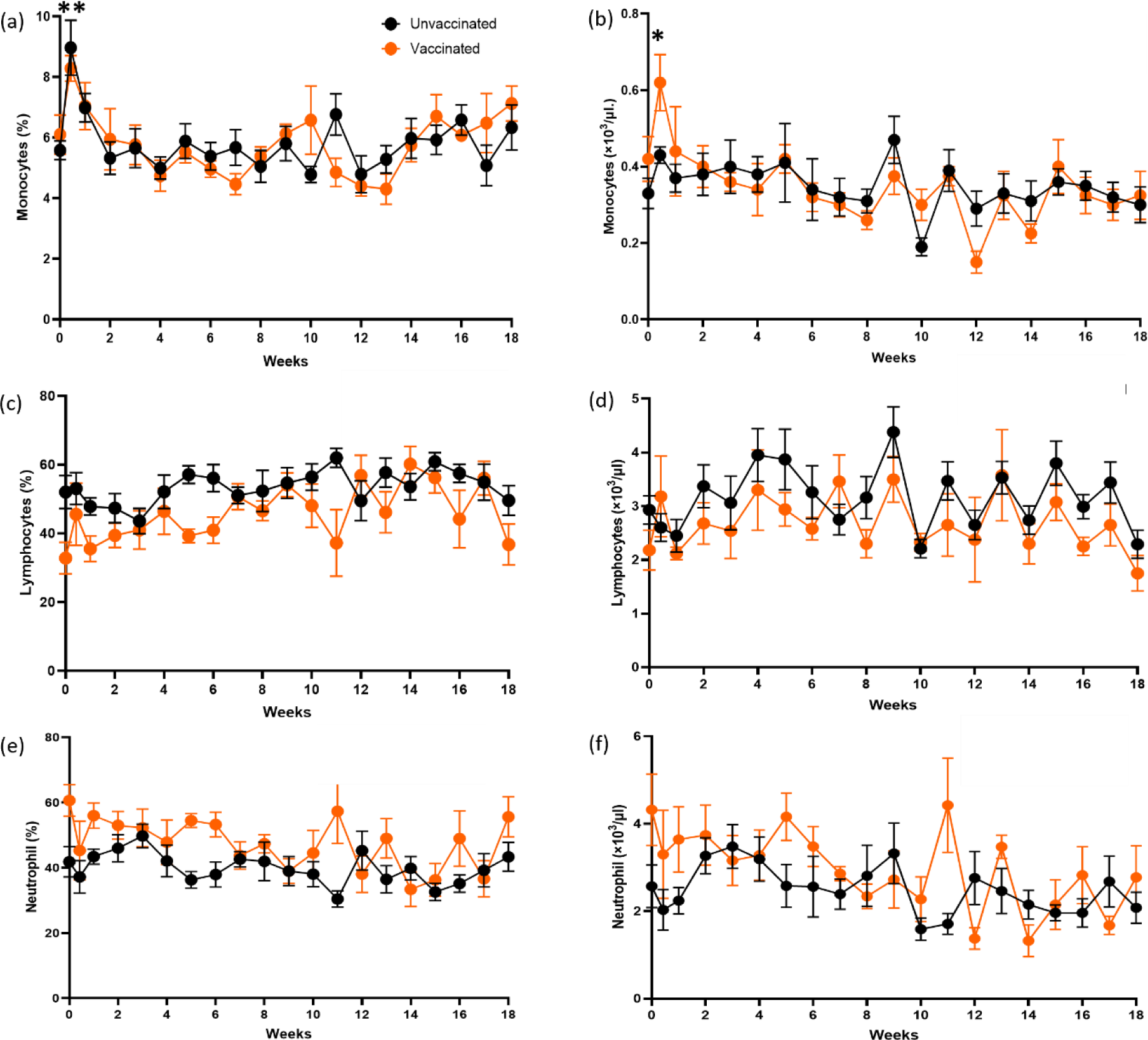
Analysis of major immune cells in blood of infected AGMs over time. (a-b) Changes in the percentage and absolute counts of monocytes, (c-d) lymphocytes, and (e-f) neutrophils in SARS-CoV-2infected vaccinated or unvaccinated AGMs. *p<0.05; **p<0.01.

**Figure S2.**
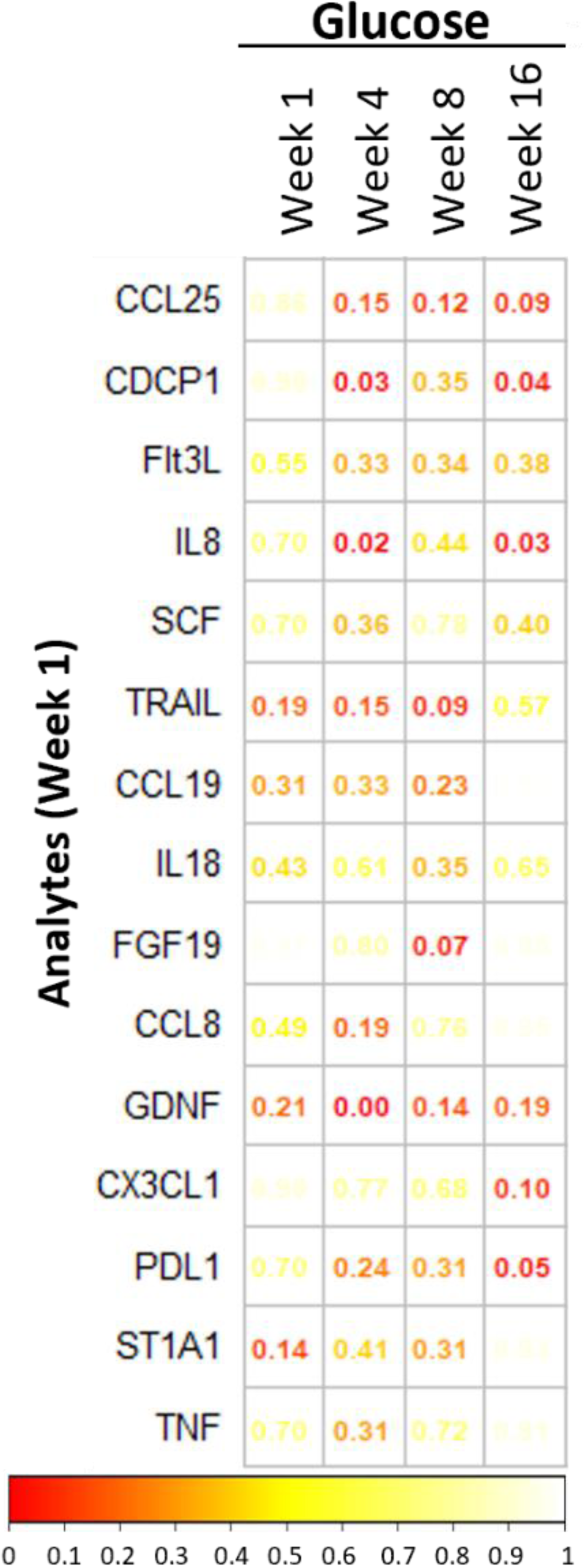
Correlation analysis between blood glucose and analytes regulated at week 1 in unvaccinated animals. Correlation matrix depicting the correlations (p values) between SARS-CoV-2-modulated analytes at week 1, and plasma glucose concentrations at specific timepoints. Spearman’s rank correlation was used for statistical analysis.

**Figure S3.**
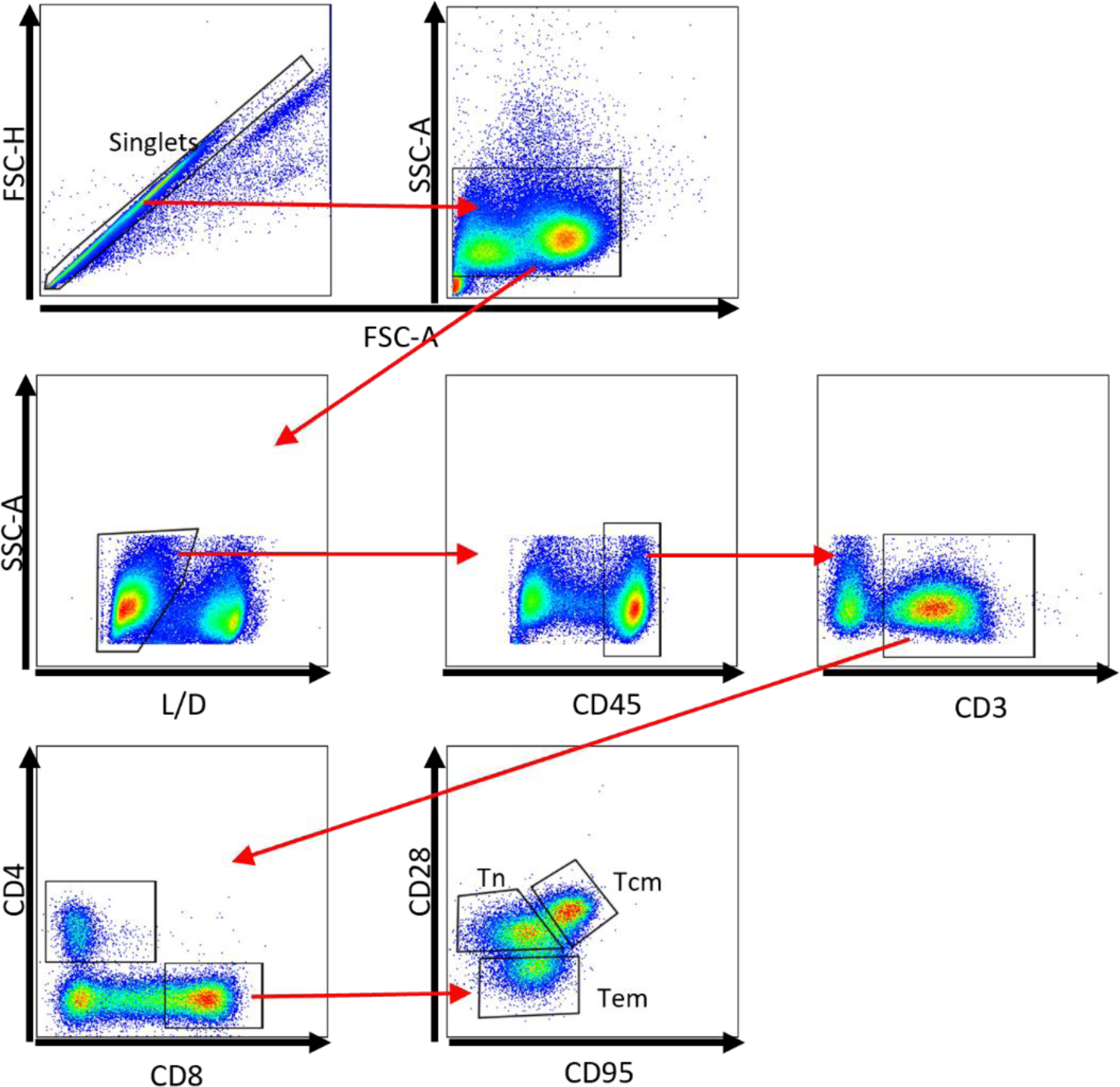
Gating strategy for Spectre and T cell hypersensitivity analysis. Red arrows show the gating steps. Tn = T naive; Tcm = T central memory; Tem = T effector memory. Total memory was defined as Tcm + Tem.

**Figure S4.**
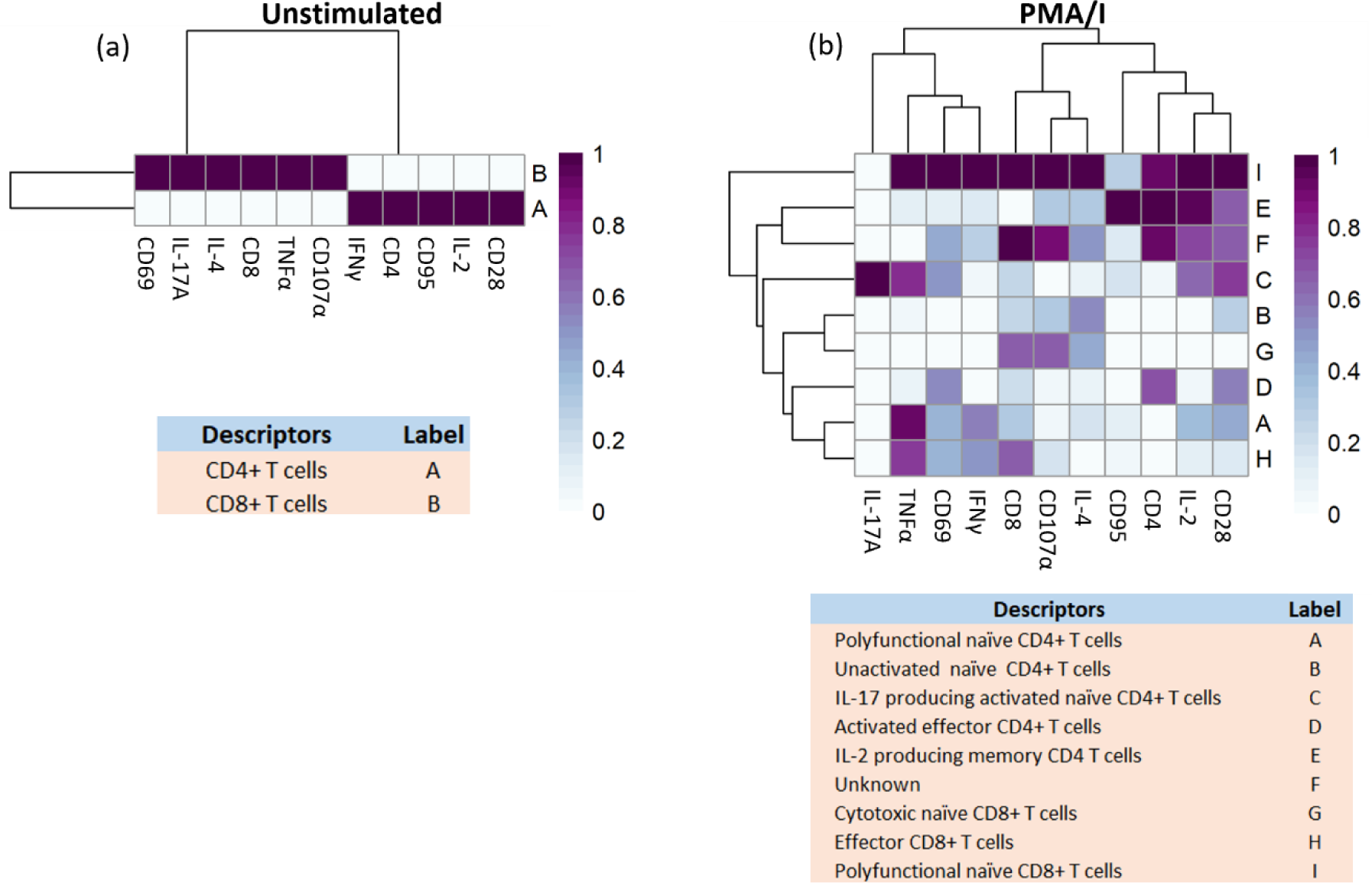
Heat maps from Spectre analysis show populations in untreated PBMCs and PMA/I treated cells. The heatmaps used to analyse and identify the unstimulated (a) and PMA/I (b) cell populations presented in Fig. 5a and 5b, respectively. CD4+ T cells are also defined as CD3+CD8-cells due to down regulation of CD4 upon PMA/I activation.

**Figure S5.**
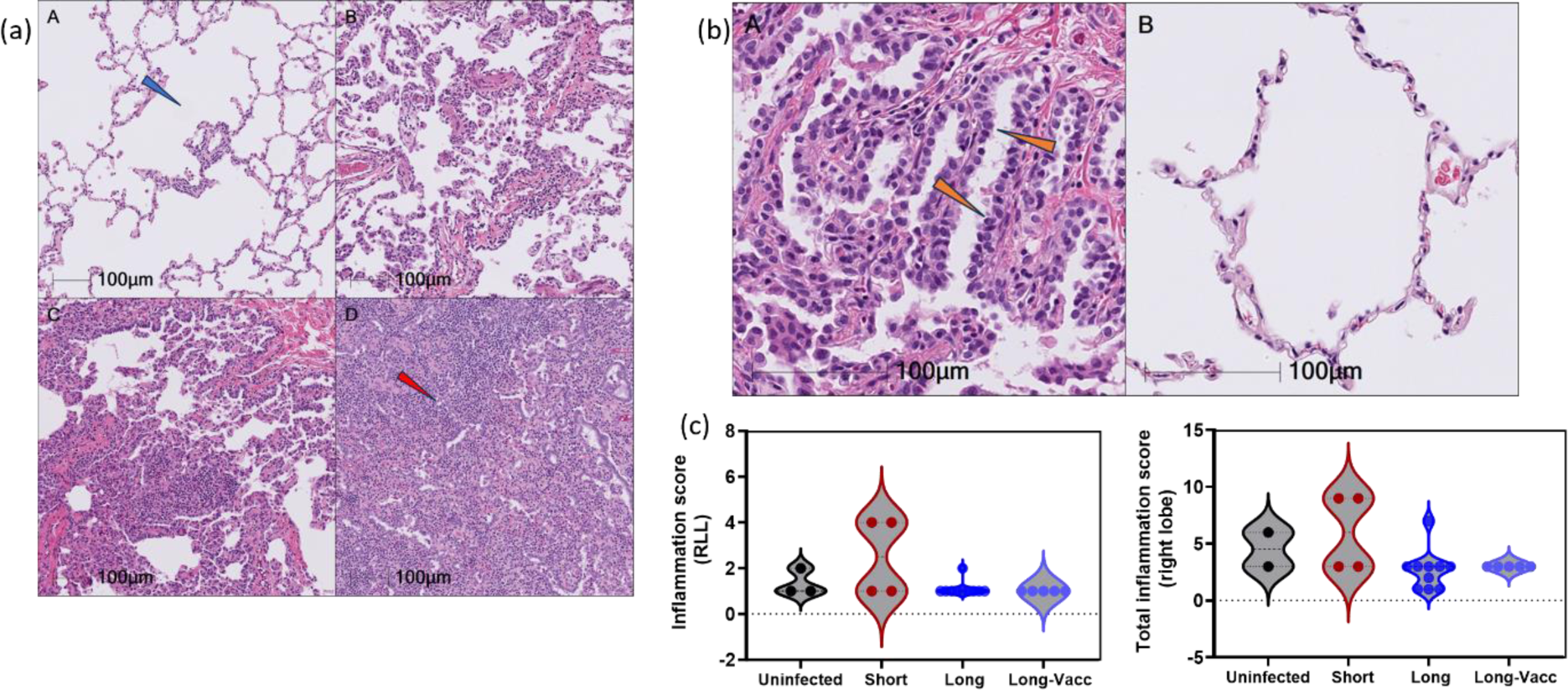
Histopathology of left anterior lungs showing representation of scoring scale used for analysis. Lung histopathology, hematoxylin & eosin stain. The histological changes in the lung were scored on the scale of 0 to 4. The changes evaluated for scoring include inflammation, pneumocyte type II hyperplasia, and fibrous connective tissue formation. (a) shows representative pictures of the histological changes and the scores. Panel A Score 1; minimal changes. Blue arrow indicates expanded alveolar spaces. Panel B. Score 2; mild changes. Panel C. Score 3; moderate changes. Panel D. Score 4; marked changes. Red arrow indicates diminished alveolar spaces due to inflammation. (b) Panel A shows a representative picture of pneumocyte type II hyperplasia from an animal infected with SARS-CoV-2. Orange arrows show marked pneumocyte type II hyperplasia. Panel B shows relatively normal alveolar septa with no pneumocyte type II hyperplasia for comparison. (c) Inflammation score of the right lower lungs (RLL, left), and composite inflammation score of the RLL (right). The scores consider the numbers of noticeable inflammatory cells present, pneumocyte type II hyperplasia, and the degree of fibrous connective tissue formation. A. Score 1; minimal. B. Score 2; mild. C. Score 3; moderate. D. Score 4; marked.

**Figure S6.**
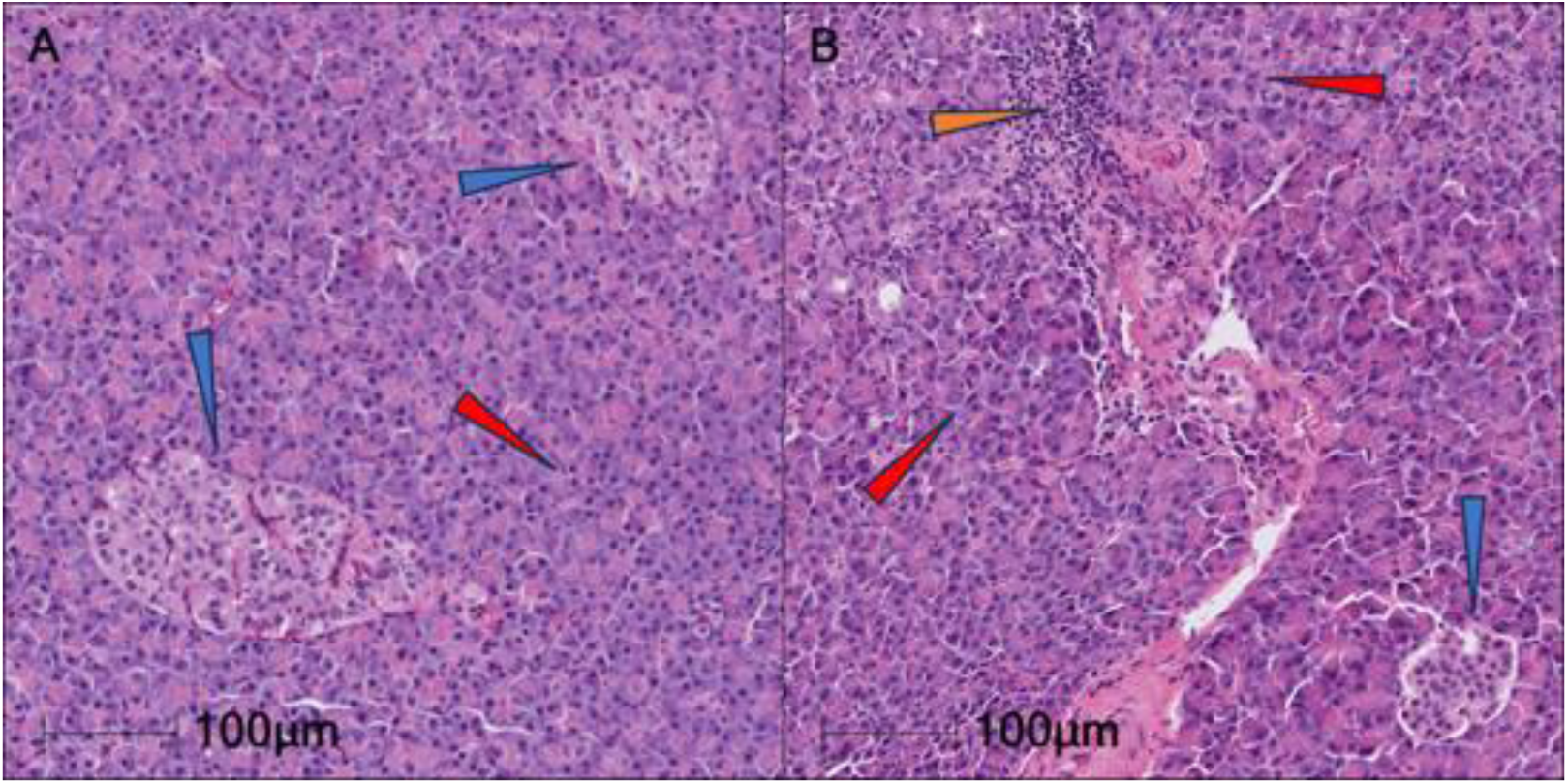
Representative H&E images of pancreas from SARS-CoV-2 infected AGMs. (A) is given an inflammation Score 0, no visible inflammation. The Islets of Langerhans (blue arrows), and exocrine pancreas (red arrow) are within normal limits. (B) is given a Score 1, minimal inflammation. The blue arrow shows an islet of Langerhans with no significant histological changes. Inflammation (orange arrow) is often within the exocrine pancreas, and periductular tissue (not shown).

**Table S1.** Significantly regulated plasma analytes between baseline and week 1 in SARS-CoV-2 infected AGMs.

| Regulated between BL and Week 1 |  |  |
| --- | --- | --- |
| Analytes | p | BH (FDR) |
| CCL25 | 0.003906 | 0.047656 |
| CDCP1 | 0.003906 | 0.047656 |
| Flt3L | 0.003906 | 0.047656 |
| MCP-2 (CCL8) | 0.003906 | 0.047656 |
| SCF | 0.003906 | 0.047656 |
| TRAIL | 0.007813 | 0.079427 |
| CCL19 | 0.011719 | 0.089355 |
| IL18 | 0.011719 | 0.089355 |
| FGF-19 | 0.019531 | 0.10831 |
| GDNF | 0.019531 | 0.10831 |
| IL8 | 0.019531 | 0.10831 |
| CX3CL1 | 0.027344 | 0.111198 |
| PD-L1 | 0.027344 | 0.111198 |
| ST1A1 | 0.027344 | 0.111198 |
| TNF | 0.027344 | 0.111198 |

**Table S2.**
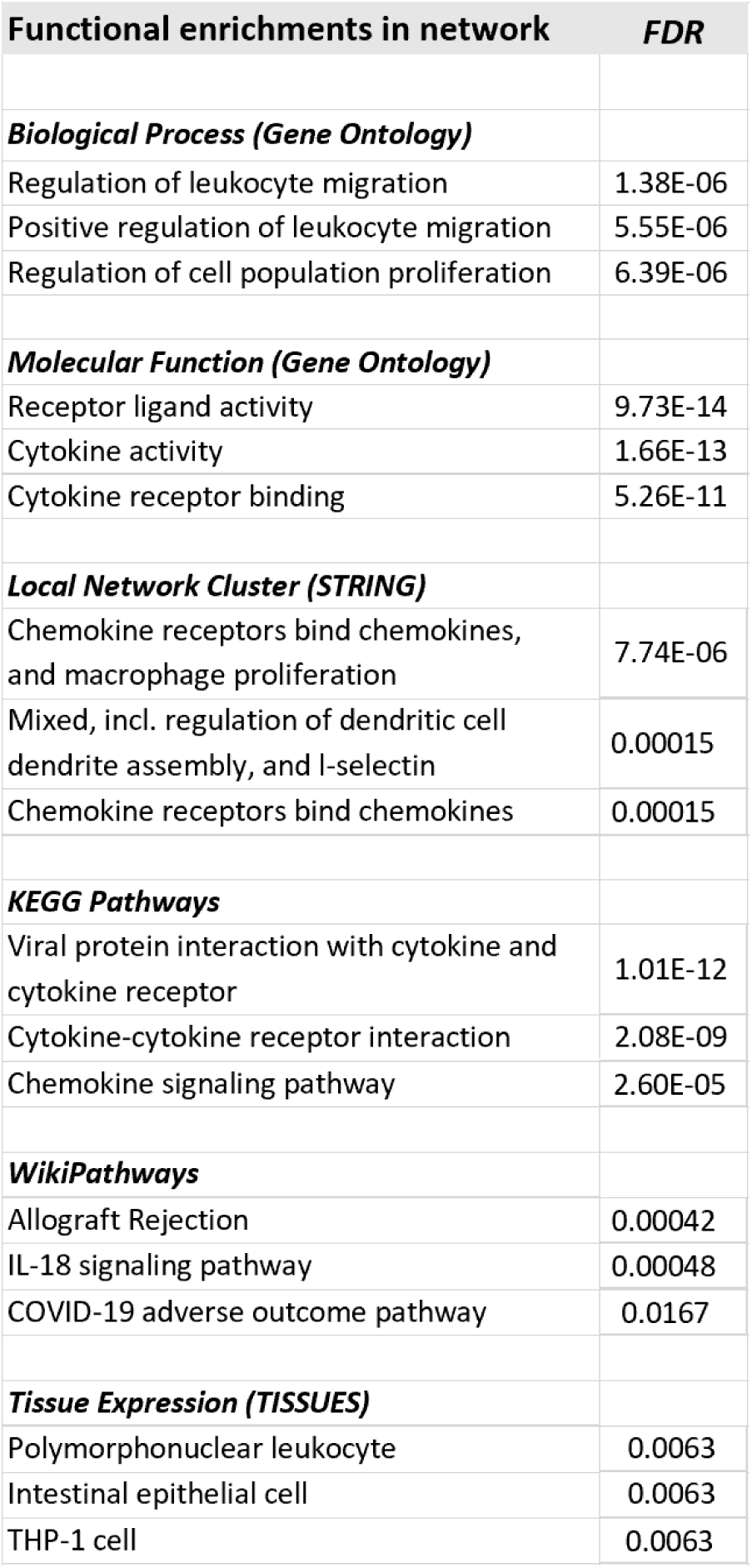
Significantly functionally enriched networks based on significantly regulated plasma analytes between baseline and week 1 in SARS-CoV-2 infected AGMs.

**Table S3.**
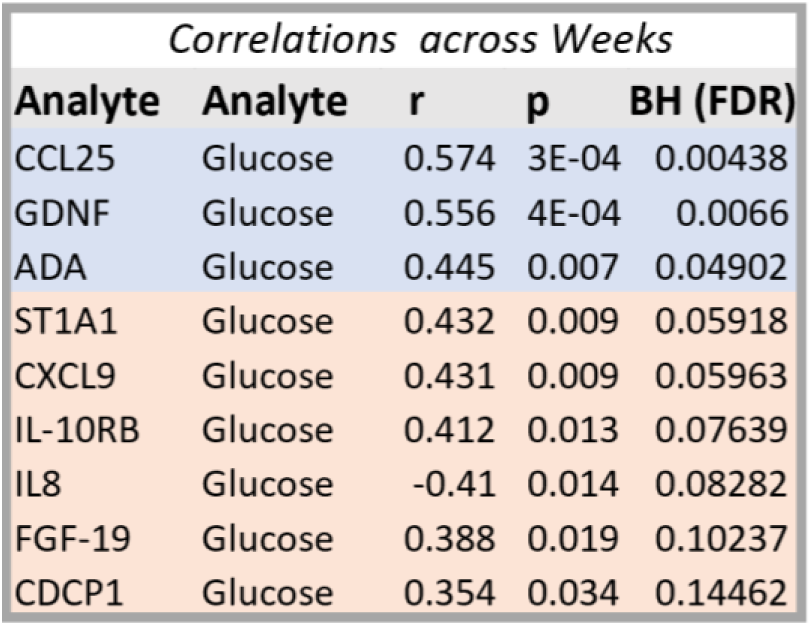
Spearman correlation analysis between blood glucose (week 0, 1, 4, 12) and plasma analytes (Olink data) in unvaccinated SARS-CoV2-infected AGMs.

## Notes

### Competing Interest Statement

The authors have declared no competing interest.

## References

1 Nalbandian, A. et al. Post-acute COVID-19 syndrome. Nat Med 27, 601–615, doi:10.1038/s41591-021-01283-z (2021).

2 Blomberg, B. et al. Long COVID in a prospective cohort of home-isolated patients. Nat Med 27, 1607–1613, doi:10.1038/s41591-021-01433-3 (2021).

3 Montefusco, L. et al. Acute and long-term disruption of glycometabolic control after SARS-CoV-2 infection. Nat Metab 3, 774–785, doi:10.1038/s42255-021-00407-6 (2021).

4 Groff, D. et al. Short-term and Long-term Rates of Postacute Sequelae of SARS-CoV-2 Infection: A Systematic Review. JAMA Netw Open 4, e2128568, doi:10.1001/jamanetworkopen.2021.28568 (2021).

5 Su, Y. et al. Multiple early factors anticipate post-acute COVID-19 sequelae. Cell 185, 881–895.e820, doi:10.1016/j.cell.2022.01.014 (2022).

6 Tang, S. W., Leonard, B. E. & Helmeste, D. M. Long COVID, neuropsychiatric disorders, psychotropics, present and future. Acta Neuropsychiatr, 1–18, doi:10.1017/neu.2022.6 (2022).

7 Mehandru, S. & Merad, M. Pathological sequelae of long-haul COVID. Nat Immunol 23, 194–202, doi:10.1038/s41590-021-01104-y (2022).

8 Wulf Hanson, S., et al. Estimated Global Proportions of Individuals With Persistent Fatigue, Cognitive, and Respiratory Symptom Clusters Following Symptomatic COVID-19 in 2020 and 2021. JAMA, doi:10.1001/jama.2022.18931 (2022).

9 Hartung, T. J. et al. Fatigue and cognitive impairment after COVID-19: A prospective multicentre study. EClinicalMedicine 53, 101651, doi:10.1016/j.eclinm.2022.101651 (2022).

10 López-Hernández, Y. et al. The plasma metabolome of long COVID patients two years after infection. Scientific reports 13, 12420, doi:10.1038/s41598-023-39049-x (2023).

11 Xie, Y. & Al-Aly, Z. Risks and burdens of incident diabetes in long COVID: a cohort study. Lancet Diabetes Endocrinol 10, 311–321, doi:10.1016/s2213-8587(22)00044-4 (2022).

12 Wang, F. et al. The laboratory tests and host immunity of COVID-19 patients with different severity of illness. JCI insight 5, doi:10.1172/jci.insight.137799 (2020).

13 Milic, J. et al. Metabolic-Associated Fatty Liver Disease Is Highly Prevalent in the Postacute COVID Syndrome. Open Forum Infectious Diseases 9, doi:10.1093/ofid/ofac003 (2022).

14 Rinaldi, R. et al. Myocardial Injury Portends a Higher Risk of Mortality and Long-Term Cardiovascular Sequelae after Hospital Discharge in COVID-19 Survivors. J Clin Med 11, doi:10.3390/jcm11195964 (2022).

15 Wu, X. et al. The Roles of CCR9/CCL25 in Inflammation and Inflammation-Associated Diseases. Front Cell Dev Biol 9, 686548, doi:10.3389/fcell.2021.686548 (2021).

16 Palmer, C. S. Innate metabolic responses against viral infections. Nat Metab 4, 1245–1259, doi:10.1038/s42255-022-00652-3 (2022).

17 Wang, W. et al. Elevated glucose level leads to rapid COVID-19 progression and high fatality. BMC Pulm Med 21, 64, doi:10.1186/s12890-021-01413-w (2021).

18 Wu, J. et al. Elevation of blood glucose level predicts worse outcomes in hospitalized patients with COVID-19: a retrospective cohort study. BMJ Open Diabetes Research & Care 8, e001476, doi:10.1136/bmjdrc-2020-001476 (2020).

19 Rahmati, M. et al. The global impact of COVID-19 pandemic on the incidence of pediatric new-onset type 1 diabetes and ketoacidosis: A systematic review and meta-analysis. J. Med. Virol. 94, 5112–5127, doi:10.1002/jmv.27996 (2022).

20 Barreto, E. A. et al. COVID-19-related hyperglycemia is associated with infection of hepatocytes and stimulation of gluconeogenesis. Proc. Natl. Acad. Sci. U. S. A. 120, e2217119120, doi:10.1073/pnas.2217119120 (2023).

21 Thorens, B. GLUT2, glucose sensing and glucose homeostasis. Diabetologia 58, 221–232, doi:10.1007/s00125-014-3451-1 (2015).

22 Han, H. S., Kang, G., Kim, J. S., Choi, B. H. & Koo, S. H. Regulation of glucose metabolism from a liver-centric perspective. Exp. Mol. Med. 48, e218, doi:10.1038/emm.2015.122 (2016).

23 Mayneris-Perxachs, J., Moreno-Navarrete, J. M. & Fernández-Real, J. M. The role of iron in host-microbiota crosstalk and its effects on systemic glucose metabolism. Nat Rev Endocrinol 18, 683–698, doi:10.1038/s41574-022-00721-3 (2022).

24 Shaw, R. J. et al. The kinase LKB1 mediates glucose homeostasis in liver and therapeutic effects of metformin. Science 310, 1642–1646, doi:10.1126/science.1120781 (2005).

25 Blair, R. V. et al. Acute Respiratory Distress in Aged, SARS-CoV-2-Infected African Green Monkeys but Not Rhesus Macaques. Am. J. Pathol. 191, 274–282, doi:10.1016/j.ajpath.2020.10.016 (2021).

26 Müller, J. A. et al. SARS-CoV-2 infects and replicates in cells of the human endocrine and exocrine pancreas. Nat Metab 3, 149–165, doi:10.1038/s42255-021-00347-1 (2021).

27 Wu, C. T. et al. SARS-CoV-2 infects human pancreatic β cells and elicits β cell impairment. Cell Metab 33, 1565–1576.e1565, doi:10.1016/j.cmet.2021.05.013 (2021).

28 Montemari, A. L. et al. An inflammatory Signature of Glucose Impairment in Cystic Fibrosis. J Inflamm Res 15, 5677–5685, doi:10.2147/jir.S365772 (2022).

29 Atanes, P., Lee, V., Huang, G. C. & Persaud, S. J. The role of the CCL25-CCR9 axis in beta-cell function: potential for therapeutic intervention in type 2 diabetes. Metabolism. 113, 154394, doi:10.1016/j.metabol.2020.154394 (2020).

30 Donath, M. Y. & Shoelson, S. E. Type 2 diabetes as an inflammatory disease. Nature Reviews Immunology 11, 98–107, doi:10.1038/nri2925 (2011).

31 Tsalamandris, S. et al. The Role of Inflammation in Diabetes: Current Concepts and Future Perspectives. Eur Cardiol 14, 50–59, doi:10.15420/ecr.2018.33.1 (2019).

32 Girard, D. & Vandiedonck, C. How dysregulation of the immune system promotes diabetes mellitus and cardiovascular risk complications. Front Cardiovasc Med 9, 991716, doi:10.3389/fcvm.2022.991716 (2022).

33 Liao, B. et al. Detection of Anti-SARS-CoV-2-S2 IgG Is More Sensitive Than Anti-RBD IgG in Identifying Asymptomatic COVID-19 Patients. Front Immunol 12, 724763, doi:10.3389/fimmu.2021.724763 (2021).

34 Qadir, M. M. F. et al. SARS-CoV-2 infection of the pancreas promotes thrombofibrosis and is associated with new-onset diabetes. JCI insight 6, doi:10.1172/jci.insight.151551 (2021).

35 Wan, L. et al. GP73 is a glucogenic hormone contributing to SARS-CoV-2-induced hyperglycemia. Nat Metab 4, 29–43, doi:10.1038/s42255-021-00508-2 (2022).

36 Liddie, S., Goody, R. J., Valles, R. & Lawrence, M. S. Clinical chemistry and hematology values in a Caribbean population of African green monkeys. J. Med. Primatol. 39, 389–398, doi:10.1111/j.1600-0684.2010.00422.x (2010).

37 Surendar, J., Mohan, V., Rao, M. M., Babu, S. & Aravindhan, V. Increased levels of both Th1 and Th2 cytokines in subjects with metabolic syndrome (CURES-103). Diabetes Technol Ther 13, 477–482, doi:10.1089/dia.2010.0178 (2011).

38 Ye, Y., Tang, X., Sun, Z. & Chen, S. Upregulated WDR26 serves as a scaffold to coordinate PI3K/ AKT pathway-driven breast cancer cell growth, migration, and invasion. Oncotarget 7, 17854–17869, doi:10.18632/oncotarget.7439 (2016).

39 Sun, Z., Tang, X., Lin, F. & Chen, S. The WD40 Repeat Protein WDR26 Binds Gβγ and Promotes Gβγ-dependent Signal Transduction and Leukocyte Migration*. Journal of Biological Chemistry 286, 43902–43912, doi:10.1074/jbc.M111.301382 (2011).

40 Penrice-Randal, R. et al. Blood gene expression predicts intensive care unit admission in hospitalised patients with COVID-19. Front Immunol 13, doi:10.3389/fimmu.2022.988685 (2022).

41 Stein, S. R. et al. SARS-CoV-2 infection and persistence in the human body and brain at autopsy. Nature 612, 758–763, doi:10.1038/s41586-022-05542-y (2022).

42 Radziuk, J. & Pye, S. Hepatic glucose uptake, gluconeogenesis and the regulation of glycogen synthesis. Diabetes. Metab. Res. Rev. 17, 250–272, doi:10.1002/dmrr.217 (2001).

43 Fahlberg, M. D. et al. Cellular events of acute, resolving or progressive COVID-19 in SARS-CoV-2 infected non-human primates. Nat Commun 11, 6078, doi:10.1038/s41467-020-19967-4 (2020).

44 Codo, A. C. et al. Elevated Glucose Levels Favor SARS-CoV-2 Infection and Monocyte Response through a HIF-1α/Glycolysis-Dependent Axis. Cell Metab 32, 437–446.e435, doi:10.1016/j.cmet.2020.07.007 (2020).

45 Lucas, C. et al. Longitudinal analyses reveal immunological misfiring in severe COVID-19. Nature 584, 463–469, doi:10.1038/s41586-020-2588-y (2020).

46 Su, Y. et al. Multi-Omics Resolves a Sharp Disease-State Shift between Mild and Moderate COVID-19. Cell 183, 1479–1495.e1420, doi:10.1016/j.cell.2020.10.037 (2020).

47 Hoel, H. et al. Elevated markers of gut leakage and inflammasome activation in COVID-19 patients with cardiac involvement. J. Intern. Med. 289, 523–531, doi:10.1111/joim.13178 (2021).

48 Ellinghaus, D. et al. Genomewide Association Study of Severe Covid-19 with Respiratory Failure. N Engl J Med 383, 1522–1534, doi:10.1056/NEJMoa2020283 (2020).

49 Muri, J., et al. Anti-chemokine antibodies after SARS-CoV-2 infection correlate with favorable disease course. bioRxiv, doi:10.1101/2022.05.23.493121 (2022).

50 Amiya, T. et al. C-C motif chemokine receptor 9 regulates obesity-induced insulin resistance via inflammation of the small intestine in mice. Diabetologia 64, 603–617, doi:10.1007/s00125-020-05349-4 (2021).

51 Chung, H. K. et al. Growth differentiation factor 15 is a myomitokine governing systemic energy homeostasis. J. Cell Biol. 216, 149–165, doi:10.1083/jcb.201607110 (2017).

52 Kim, K. H. et al. Growth differentiation factor 15 ameliorates nonalcoholic steatohepatitis and related metabolic disorders in mice. Scientific reports 8, 6789, doi:10.1038/s41598-018-25098-0 (2018).

53 Mullican, S. E. et al. GFRAL is the receptor for GDF15 and the ligand promotes weight loss in mice and nonhuman primates. Nat Med 23, 1150–1157, doi:10.1038/nm.4392 (2017).

54 Emmerson, P. J. et al. The metabolic effects of GDF15 are mediated by the orphan receptor GFRAL. Nat Med 23, 1215–1219, doi:10.1038/nm.4393 (2017).

55 Parchwani, D., Dholariya, S., Katoch, C. & Singh, R. Growth differentiation factor 15 as an emerging novel biomarker in SARS-CoV-2 infection. World J Methodol 12, 438–447, doi:10.5662/wjm.v12.i5.438 (2022).

56 Anitha, M. et al. GDNF rescues hyperglycemia-induced diabetic enteric neuropathy through activation of the PI3K/Akt pathway. J Clin Invest 116, 344–356, doi:10.1172/jci26295 (2006).

57 Beg, M., Abdullah, N., Thowfeik, F. S., Altorki, N. K. & McGraw, T. E. Distinct Akt phosphorylation states are required for insulin regulated Glut4 and Glut1-mediated glucose uptake. eLife 6, doi:10.7554/eLife.26896 (2017).

58 Zisman, A. et al. Targeted disruption of the glucose transporter 4 selectively in muscle causes insulin resistance and glucose intolerance. Nat Med 6, 924–928, doi:10.1038/78693 (2000).

59 Aldhshan, M. S., Jhanji, G., Poritsanos, N. J. & Mizuno, T. M. Glucose Stimulates Glial Cell Line-Derived Neurotrophic Factor Gene Expression in Microglia through a GLUT5-Independent Mechanism. International journal of molecular sciences 23, doi:10.3390/ijms23137073 (2022).

60 Tang, M. J., Worley, D., Sanicola, M. & Dressler, G. R. The RET-glial cell-derived neurotrophic factor (GDNF) pathway stimulates migration and chemoattraction of epithelial cells. J. Cell Biol. 142, 1337–1345, doi:10.1083/jcb.142.5.1337 (1998).

61 Young, H. M. et al. GDNF is a chemoattractant for enteric neural cells. Dev. Biol. 229, 503–516, doi:10.1006/dbio.2000.0100 (2001).

62 Yang, Y. et al. THE ASSOCIATION OF DECREASED SERUM GDNF LEVEL WITH HYPERGLYCEMIA AND DEPRESSION IN TYPE 2 DIABETES MELLITUS. Endocr Pract 25, 951–965, doi:10.4158/ep-2018-0492 (2019).

63 Mwangi, S. et al. Glial cell line-derived neurotrophic factor increases beta-cell mass and improves glucose tolerance. Gastroenterology 134, 727–737, doi:10.1053/j.gastro.2007.12.033 (2008).

64 Gebre, M. S. et al. mRNA vaccines induce rapid antibody responses in mice. NPJ Vaccines 7, 88, doi:10.1038/s41541-022-00511-y (2022).

65 Taylor, K. et al. Diabetes following SARS-CoV-2 infection: Incidence, persistence, and implications of COVID-19 vaccination. A cohort study of fifteen million people. medRxiv, 2023.2008.2007.23293778, doi:10.1101/2023.08.07.23293778 (2023).

66 Trypsteen, W., Van Cleemput, J., Snippenberg, W. V., Gerlo, S. & Vandekerckhove, L. On the whereabouts of SARS-CoV-2 in the human body: A systematic review. PLoS Pathog 16, e1009037, doi:10.1371/journal.ppat.1009037 (2020).

67 Fears, A. C. et al. Exposure modality influences viral kinetics but not respiratory outcome of COVID-19 in multiple nonhuman primate species. PLoS Pathog 18, e1010618, doi:10.1371/journal.ppat.1010618 (2022).

68 Beddingfield, B. J. et al. Effective Prophylaxis of COVID-19 in Rhesus Macaques Using a Combination of Two Parenterally-Administered SARS-CoV-2 Neutralizing Antibodies. Front Cell Infect Microbiol 11, 753444, doi:10.3389/fcimb.2021.753444 (2021).

69 Arunachalam, P. S. et al. Durable protection against the SARS-CoV-2 Omicron variant is induced by an adjuvanted subunit vaccine. Science translational medicine 14, eabq4130, doi:10.1126/scitranslmed.abq4130 (2022).

70 Ashhurst, T. M. et al. Integration, exploration, and analysis of high-dimensional single-cell cytometry data using Spectre. Cytometry A 101, 237–253, doi:10.1002/cyto.a.24350 (2022).

71 R: A language and environment for statistical computing. R Foundation for Statistical Computing (2022).

72 Assarsson, E. et al. Homogenous 96-plex PEA immunoassay exhibiting high sensitivity, specificity, and excellent scalability. PLoS One 9, e95192, doi:10.1371/journal.pone.0095192 (2014).

73 Schindelin, J. et al. Fiji: an open-source platform for biological-image analysis. Nat Methods 9, 676–682, doi:10.1038/nmeth.2019 (2012).

74 Szklarczyk, D. et al. STRING v10: protein-protein interaction networks, integrated over the tree of life. Nucleic Acids Res 43, D447–452, doi:10.1093/nar/gku1003 (2015).

75 Mi, H., Muruganujan, A. & Thomas, P. D. PANTHER in 2013: modeling the evolution of gene function, and other gene attributes, in the context of phylogenetic trees. Nucleic Acids Res 41, D377–386, doi:10.1093/nar/gks1118 (2013).

76 Thomas, P. D. et al. PANTHER: Making genome-scale phylogenetics accessible to all. Protein Sci. 31, 8–22, doi:10.1002/pro.4218 (2022).

